# Human settlement drives population divergence of *Leishmania guyanensis* at the sylvatic-anthropogenic interface

**DOI:** 10.64898/2026.03.06.710114

**Authors:** João Luís Reis-Cunha, Luiza C. Reis, Luciana Mendes, Melissa Cavalcante, Mathew Johansson, Colin J. McClean, Maria das Graças Vale Barbosa Guerra, Hiro Goto, Jorge Augusto de Oliveira Guerra, Daniel C. Jeffares

## Abstract

The Amazon is the largest tropical rainforest in the world, and is an important interface between natural rainforest and anthropogenic environments. In the central Amazon region, *Leishmania guyanensis* is a common agent of cutaneous leishmaniasis (CL), a disfiguring vector-borne disease that circulates between wild mammalian reservoirs and sandfly vectors in sylvatic cycles. People become infected when they interact or disturb this cycle. At present, little is known about the adaptation of *Leishmania* parasites at the interface between sylvatic and anthropogenic environments. Here, to characterize CL in a transition zone, we conducted a population genomic analysis of 77 *Leishmania* isolates from human patients living in Manaus’s outskirts or in highways adjacent to the Brazilian Amazon rainforest. We discovered two *L. guyanensis* populations; an ancestral ‘sylvatic’ population occurring in densely forested areas far from the city and a derived population, occurring in areas with lower forest cover closer to the city (the ‘city-proximal’ population). The genomic divergence between these populations indicates that their separation occurred relatively recently, consistent with human-driven development of a new transmission cycle for CL in the Manaus region. The lack of recent gene flow between these populations indicates that the newer city-proximal population has established a transmission cycle that operates independently from ancestral sylvatic transmission. This provides a striking example of the adaptation of disease transmission to an environment that has been altered by human activity.

**Significance Statement:** Emerging infectious diseases often arise at the interface between natural and human-modified environments. One of these important interfaces is the Amazon, the largest rainforest in the world. We analyzed the population genomics of *Leishmania guyanensis*, the agent of cutaneous leishmaniasis in the Amazon, typically transmitted in sylvatic cycles. Our findings reveal a recent emergence of a *L. guyanensis* population associated with anthropogenic environment alterations, which is genetically distinct from the ancestral sylvatic lineage. This newly adapted population highlights how landscape change and urban expansion may drive the evolution and emergence of protozoan vector-borne pathogens at the rainforest–urban interface.

## Introduction

The Amazon is the largest tropical rainforest in the world, covering 35-40% of the South America continent and extending over 9 countries, with ∼60% of its extension in Brazil (1–3). It is an important interface between rainforest and urban environments (4–6), as it harbours 10% of the global biodiversity, and is home to more than 30 million people. Deforestation and poverty create favourable conditions for the transmission of numerous vector-borne tropical infectious diseases in the region, such as malaria, Chagas disease, dengue, visceral and cutaneous leishmaniasis (*1, 7*). Understanding how pathogens, vectors, and reservoirs move from forest to rural areas is essential for disease control in a context of environmental management.

Cutaneous leishmaniasis (CL) is a vector-borne disease with chronic evolution and several clinical forms with increasing severity, such as: localised cutaneous leishmaniasis (LCL), diffuse cutaneous leishmaniasis (DCL), and mucocutaneous (MCL) leishmaniasis. While mortality rate for CL is low, it causes a large burden due to the disease’s chronicity, which can cause social stigma due to scars and disfigurement (8, 9). In South America, there are 180,000-300,000 cases of CL reported each year (8, 10, 11). In Brazil 40% of the 20-30,000 cases reported each year are within the Amazon region (12, 13).

CL in the Amazon is primarily a zoonosis, circulating between wild mammal reservoirs and sandfly vectors in sylvatic cycles (14–16). Human infections are a consequence of events that disturb this cycle or human activities that result in contact with infected insect vectors in forests, such as deforestation, mining, farming and livestock production (14, 17). Seven *Leishmania* species are reported as etiological agents for CL in the Brazilian Amazon: *Leishmania* (*Viannia*) *braziliensis*, *Leishmania* (*Viannia*) *guyanensis*, *Leishmania* (*Viannia*) *naiffi*, *Leishmania* (*Viannia*) *lainsoni*, *Leishmania* (*Viannia*) *shawi*, *Leishmania* (*Viannia*) *lindenbergi* and *Leishmania* (*Leishmania*) *amazonensis*, which vary in associations with vectors, reservoirs and clinical outcomes (14, 18).

*L. guyanensis* is the most prevalent cause of human CL cases in Manaus, which contrasts with other regions in South America where the majority of cases are caused by *L.* (V.) *braziliensis* (*13, 19–23*). Manaus is the largest city in western Amazon with 2.4 million people, and has an incidence of 1,000-4,000 CL cases reported per year (14). Unplanned urbanization and deforestation result in frequent outbreaks in the city outskirts or in settlements along two highways that connect the city to other regions in the Amazon (13, 14, 24). Since *L. guyanensis* can cause the severe disfiguring MCL (22), understanding its transmission routes and adaptive strategies becomes particularly important.

Despite its epidemiologic importance, relatively little is known about *L. guyanensis* eco epidemiology, genetic diversity or adaptation to different environments. The main vector of *L. guyanensis* in the Manaus region is *Nyssomyia umbratilis*, but other species such as *Lutzomyia whitmani* and *Lutzomyia anduzei* were reported harbouring this parasite in other regions (25–28). Similarly, a variety of mammalian species are known to harbor *L. guyanensis* parasites (26, 29–32). The occurrence of *L. guyanensis* in different vectors and reservoirs is likely to alter transmission dynamics in different biomes. An early population genomics study of *L. guyanensis* in French Guiana showed that parasite populations were structured with geographical distances, with evidence of frequent sexual reproduction and well delimited microfoci. Estimates of migration rates were partially explained by short distance dispersal occurring via vectors and reservoirs, rather than long distance movements, consistent with a primarily sylvatic maintenance of transmission (33).

In the present study, we analysed the genomes of 77 *Leishmania* clinical isolates from patients that resided in Manaus and neighbouring areas. We confirmed 71 as *L. guyanensis*, and demonstrate that a derived population of *L. guyanensis* is predominantly found in areas closer to Manaus with less forest coverage (the ‘proximal’ population, LguyP), which emerged recently from an ancestral sylvatic population (the ‘sylvatic’ population, LguyS) that persists in forested regions. We speculate that this separation is due to adaptation of vectors and/or reservoirs to transition zones between the city and heavy forested areas, due to anthropogenic change. This represents an early phase in development of leishmaniasis transmission cycles that operate independently of natural ecosystems, which has important epidemiological implications.

## Results

### Characterization of the *Leishmania* populations in Manaus region

The 77 evaluated *Leishmania* primary isolates were obtained from biopsies from human patients that live in the city of Manaus and on highways adjacent to the Brazilian Amazon rainforest up to 240 km from the city. These patients were admitted and treated at the Fundação de Medicina Tropical Heitor Vieira Dourado (FMTHVD) in Manaus, from September 2018 to March 2020. The parasites were briefly cultured prior to sequencing and the *Leishmania* species was initially identified by PCR (22, 34, 35).

To confirm the identity of these 77 *Leishmania* isolates, we compared their genomes with representative samples from seven *L. Viannia* species that circulate in the Amazon: *L. braziliensis*, *L. guyanensis*, *L. lainsoni*, *L. naiffi*, *L. panamensis*, *L. peruviana* and *L. shawi* (Figure 1, Supplementary Figure 1, Supplementary Table 1). From the 77 primary isolates, 71 clustered in a single branch that was close to previously characterised *L. guyanensis* samples from Brazil and Venezuela in the phylogenetic network (strains 204-365, M4147 and S8104, Figure 1A blue box, Supplementary Figure 1). Species classification was further confirmed by PCA (Supplementary Figure 1). Four of the *L. guyanensis* primary isolates (P13, P57, P59, P67) had evidence of being polyploid or multiclonal, based on deviations in the expected distribution of allelic read depths (Complexity Index, CI) (36) (Supplementary Figure 2 B and C), and by the high heterozygous/homozygous SNP ratios (P13, P57, P67) (Supplementary Figure 2 A). The six isolates that were not *L. guyanensis* were classified as other *L. Viannia* species that are known to circulate in the amazon: P15, P43 and P47 as *L. braziliensis*; P20 and P56 as *L. naiffi* and P05 as *L. lainsoni* (Figure 1 A, Supplementary Figure 1). From these, sample P43 appeared to be a polyploid hybrid or multi-species infection, with genetic contributions from *L. braziliensis*, *L. peruviana*, *L. panamensis* and *L. naiffi* populations (Figure 1 D, red arrow, Supplementary Figure 2). Two other samples, P15 and P47 are likely hybrids, with major genetic contributions from *L. braziliensis* and a minor contribution from *L. peruviana* (Figure 1 C, Supplementary Figure 2). The occurrence of chromosome duplications was rare, and the only consistent supernumerary chromosome observed was chromosome 31 (37) (Supplementary Figure 5). The genomic nucleotide diversity (π) of the 71 confirmed *L. guyanensis* primary isolates is similar to what is observed in *L. peruviana*, higher than *L. panamensis* and lower than the *L. braziliensis* datasets evaluated here (Figure 1 B).

**Figure 1:**
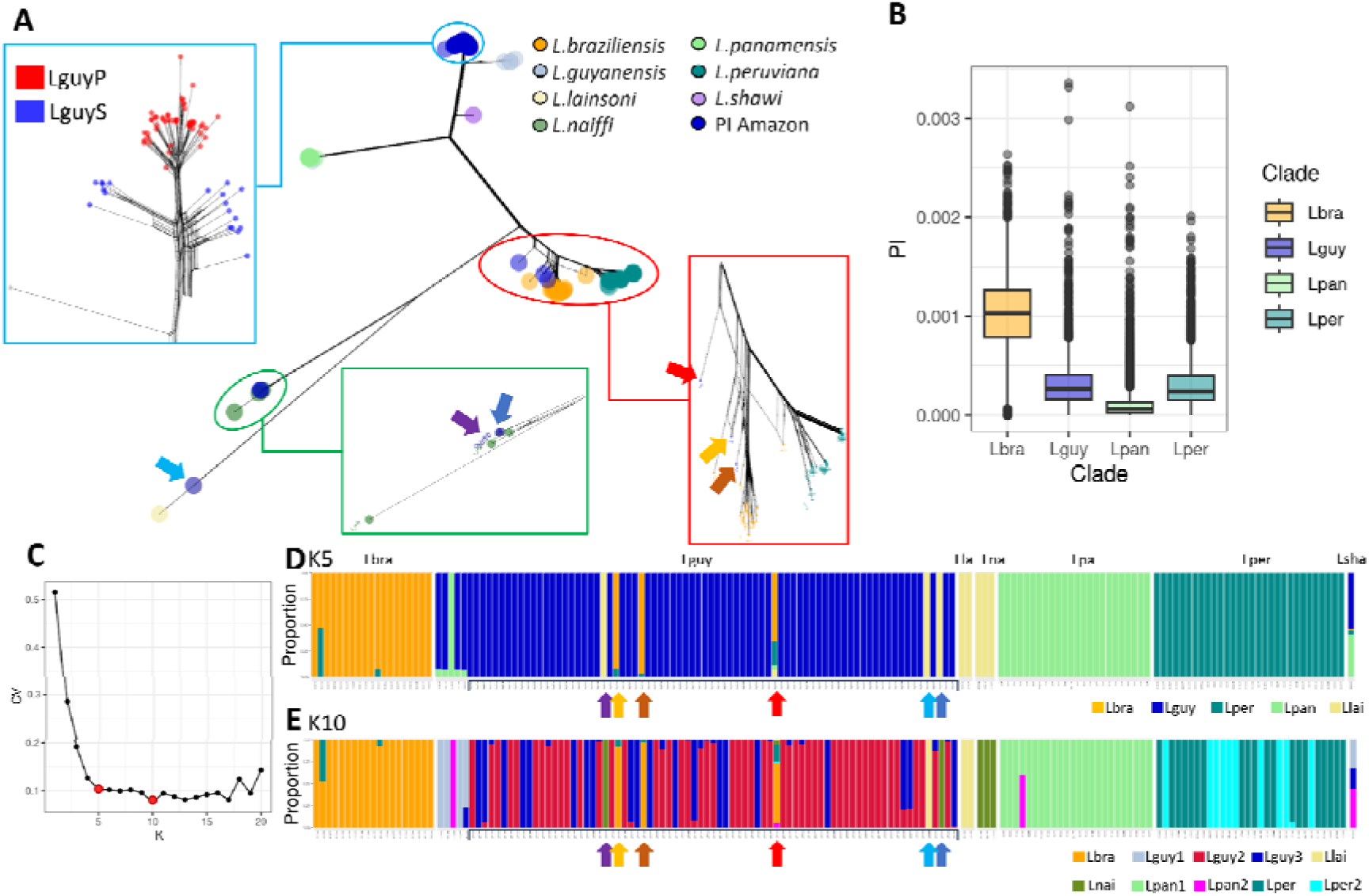
*Leishmania guyanensis* phylogeny and population structure. Comparison of the *L. guyanensis* primary isolates from Amazon (PI Amazon) and representative isolates from *Leishmania* Viannia subgenus: *L. braziliensis*, *L. guyanensis*, *L. lainsoni*, *L. naiffi*, *L. panamensis*, *L. peruviana* and *L. shawi*. The six arrows point to primary isolates that were classified as *L.* guyanensis by PCR, but correspond to other species or hybrids. **A)** Phylogenetic network based on the nuclear genome SNPs. The blue box highlights the branch with the true *L. guyanensis* isolates from Amazon, re-colouring the isolates based on their main population on the Admixture analysis with K=10 (LguyP or LguyS). The green and red boxes highlight, respectively, branches with *Leishmania* isolates from the Amazon that were classified as *L. naiffi* or *L. braziliensis*. **B)** Nucleotide diversity of the evaluated species with more than 5 samples, using 50 kb windows. The mean diversity for each are: Lbra=0.001, Lguy=0.000316; Lpan=0.000123; Lper=0.000313. **C)** Admixture cross validation error plot for each K value. K5 and K10 are highlighted by red dots. **D)** *L. Viannia* Admixture, using K=5 and K=10. In this panel, each sample is grouped based on its pre-admixture species assignment: Lbra: *L. braziliensis*; Lguy: *L. guyanensis*; Lla: *L. lainsoni*; Lna: *L. naiffi*; Lpa: *L. panamensis*; Lper:; *L. peruviana* and Lsha: *L. shawi*. The blue box highlights the Amazonian *L. guyanensis* primary isolates. The arrows represent the isolates in the phylogeny network in A.

We evaluated the *L. guyanensis* population structure using Admixture (38), with two stratification values that had low cross-validation error: K=5 and K=10 (Figure 1 C). Using K=5 we were able to confirm the 71 primary isolates as being a part of the *L. guyanensis* population (Figure 1, D), while the use of K=10 uncovered two *L. guyanensis*’ populations coexisting at the same time in the Manaus region (Figure 1 A and E). They were named Lguy Sylvatic (LguyS) and Lguy city Proximal LguyP, based on their distance to the city of Manaus (see below). These populations differ from the population of the M4147 type strain obtained in Brazil in 1945, and will be further explored in the next sections.

### The ancestral population (LguyS) is mainly observed in forested outskirts, while the derived population (LguyP) predominates in regions closer to the city with greater deforestation

Evaluation of the phylogenetic network and PCA of the *L. guyanensis* strains (Figure 1A, Figure 2A) confirmed the presence of two populations, LguyP with 51 isolates and LguyS with 20 isolates. P44 was removed from LguyS in downstream analysis, as it is an outgroup (Figure 1A and 2A), leaving 19 isolates. To understand the environment where these two populations reside, we examined the geographic locations, the distance to the city of Manaus and the forest cover around each of the patients’ homes when the *L. guyanenses* isolates were collected, from 2018–2020 (Figure 2 A-C). Most of the samples from LguyS were isolated from patients that lived in houses or settlements along the two main forest-crossing highways that leave the city of Manaus to the North and East: AM-010 and BR-174 (median distance from Manaus = 96 km). Isolates from LguyP were obtained from patients that resided significantly closer to the city of Manaus (median distance from Manaus = 35 km, Mann Whitney U test p-value = 3E^-08^) (Figure 2 A). Forest cover in a 3 km range around residences was significantly higher for residences of patients infected with LguyS parasites (median forest cover = 93%) compared with residences from patients with LguyP (median forest cover = 85%) (Mann Whitney U test, *P* = 0.0047) (Figure 2 C). Significant differences in forest cover were also obtained with radiuses of 2 to 10 km ranges around the residences (Figure 2 C, Supplementary Figure 6). Hence, the derived population (LguyP) is found closer to the city of Manaus in regions where forest cover is lower.

**Figure 2:**
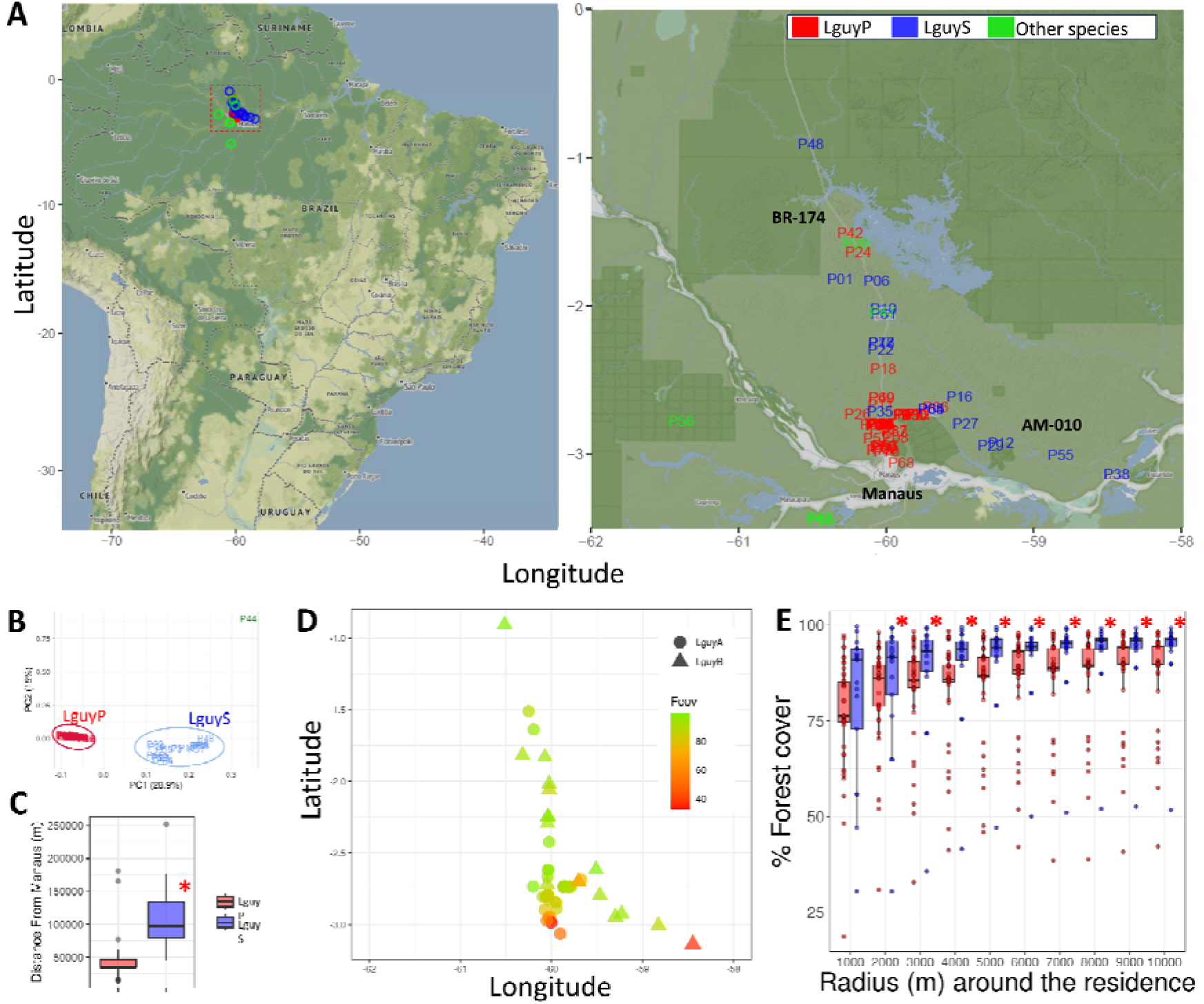
*L. guyanensis* has a distal sylvatic (LguyS) and a city-proximal (LguyP) population near Manaus. **A)** Geographic distribution of CL patient residences bearing the *L. guyanensis* primary isolates. LguyS, LguyP and other leishmania species are represented respectively in blue, red and green. **Left panel**: Continental-scale map showing the collection sites, with each isolate represented by a colored circle. **Right panel:** Expanded view of the highlighted region (red box), showing individual isolates labeled by their ID. The locations of the city of Manaus, and the main roads BR-174 and AM-010 are also shown. **B)**: Whole genome SNP PCA confirms the presence of two populations, LguyP (red) and LguyS (blue), and one outlier isolate, P44. **C):** boxplot representing the distribution of the distances of the residences from patients infected with LguyS and LguyP to the city of Manaus (Mann Whitney U test p-value = 3E-08). **D)** Percentage of forest coverage around 3 km of the local area of residence of each patient (Fcov). Other radius values can be seen in Supplementary Figure 6. **E)** Boxplot representing the percentage of forest coverage surrounding patient residences with increasing radius values (1000 to 10,000 m). Significant differences between LguyS and LguyP are represented by red asterisks.

### LguyP is derived from LguyS, and inter-population recombination events are rare

As both populations coexisted temporally and have some geographic overlap, we evaluated if there is evidence for genetic exchange between them (Figure 3 A and Supplementary Figure 8-10). LguyP-LguyS had a moderate weighted F_ST_ value of 0.158 (the proportion of the genetic variance contained within populations, compared with the total genetic variance). This is consistent with the clear differentiation of populations we observe from PCA and phylogenetic analysis (Figure 1A and 2 A). To assess the extent of recent gene flow between LguyP and LguyS, we generated individual chromosomal PCAs based on SNP data, which consistently grouped LguyP and LguyS strains in different clusters. The little evidence of inter-population genetic exchanges was restricted to a few chromosomes from few isolates (Figure 3 A and B), which was supported by analysis of population-diagnostic SNPs (Supplementary Figure 8-10). To examine this further, we used the Diagnostic Index Expectation Maximisation (*diem*) method (39) (https://cran.r-project.org/web/packages/diemr/), which assigns marker alleles to population groups while simultaneously identifying which individuals belong to each population and detecting introgressed genomic regions. *diem* identified the same populations as the PCA and Admixture analyses, with no clear evidence of recent introgression (Supplementary Figure 14). As the separation of these two populations is recent, they still share alleles across the genome, which might reduce the power of diem to identify the introgressed regions. In summary, our analyses indicate that these two populations are separating, and recent genetic exchange between these two populations is not common, consistent with two largely independent transmission cycles.

**Figure 3:**
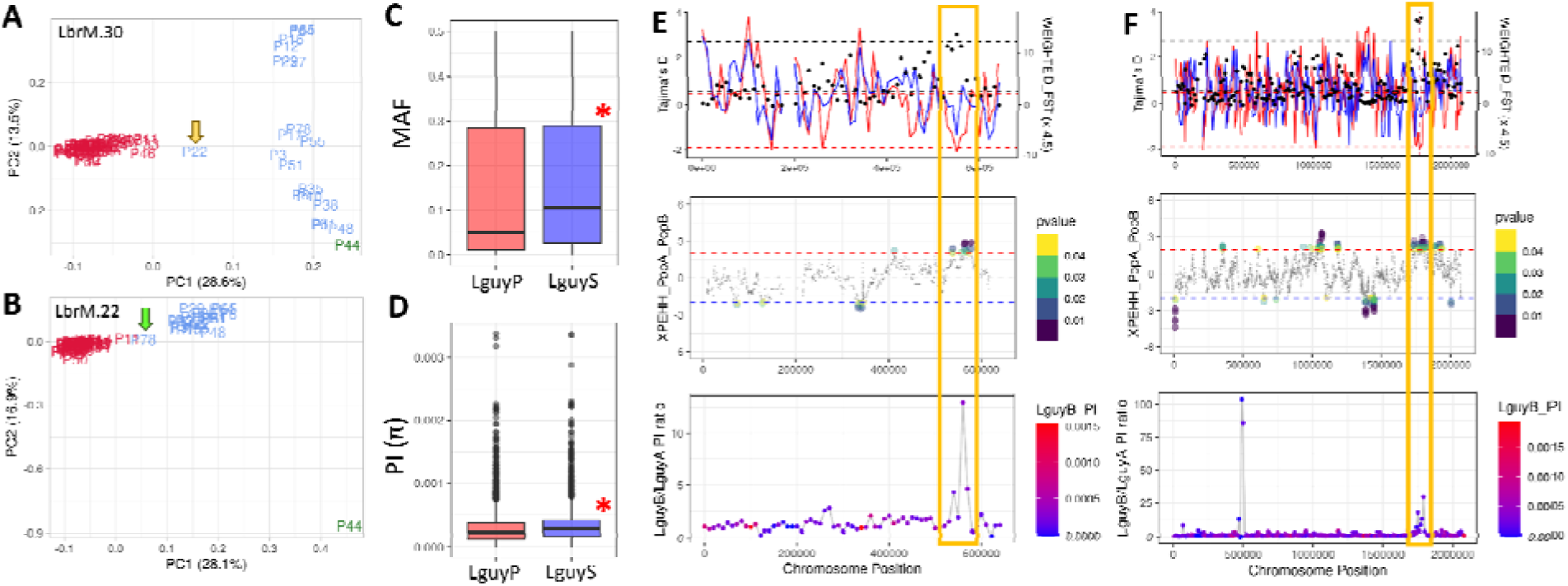
Evidence for recombination and selection in LguyP and LguyS. We observe evidence for genetic exchange between populations, in Principal component analysis (PCA) of **A)** chromosome 30 (LbrM.30) SNPs, showing the isolate P22 in an intermediary position between LguyP and LguyS, and also in **B)** chromosome 22 (LbrM.22) PCA that strain P78 in an intermediary position between LguyP and LguyS. Minor allele frequency (MAF) **C)** and nucleotide diversity (π) **D)** comparing LguyP and LguyS. Evidence for selective sweeps in two chromosomal regions: chromosome 15 **(E)** and chromosome 34 **(F)**. The top panel shows Tajima’s D (left Y axis, coloured lines, red: LguyP; blue: LguyS) and F_ST_ (right Y axis, black dots) along the chromosomal region. The mid panel corresponds to XP-EHH results (Y axis) for each SNP position along the chromosome comparing LguyP and LguyS. Each dot corresponds to one SNP, where significant p-values are larger and coloured. Values above the red line correspond to potential selective sweeps on LguyP, while values below the blue line to potential sweeps in LguyS. The bottom panel corresponds to the nucleotide diversity (π) ratio between LguyS and LguyP, coloured by nucleotide diversity (PI) in LguyS. The gold boxes highlight regions with low Tajima’s D, high F_ST_, high XP-EHH result and higher nucleotide diversity in LguyS, indicating a potential selective sweep and loss of diversity in the derived peridomestic population LguyP.

Several pieces of evidence support that LguyP is a younger population derived from the LguyS. LguyP forms a derived branch in the phylogeny network, which is external when compared to other *Leishmania* clades (Figure 1 A), a tighter cluster in PCA (Figure 2 A), and has a ∼15% lower nucleotide diversity (2.91 x 10^-3^) when compared to LguyS (3.37 x 10^-3^) in genome-wide 10 kb windows (Mann Whitney U test p-value = 1.19 x 10^-26^) (Figure 3 C and D). To examine the time scale within which the LguyP population emerged, we calculated the *d*_XY_, the mean genetic distance among all between-population strain pairs, using the kinetoplastid maxicircle (mitochondrial) DNA sequences (kDNA), which we expect to diverge in a tree-like manner (Supplementary Figure 3). Between population mean kDNA SNP counts were 1.33 (*d*_XY_ x 1000 nt = 0.081). In context, this is approximately half of what we observe between *L. infantum* populations from Brazil and Europe (36), with a mean SNP count of 3 (*d*_XY_ x 1000 nt = 0.176). As the Brazil-Europe *L. infantum* population was previously estimated to have diverged between 1648 and 1941 CE (95% HDPI) (40), the LguyP-LguyS split should be more recent.

To utilise the entire nuclear genomic variation, we examined the genome-wide population divergence, using F_ST_. In our analysis using a much larger set of *Leishmania donovani* species complex genomes, nuclear genomic divergence showed that population divergence F_ST_ was correlated with population divergence times, until F_ST_ was saturated at ∼0.6 (Supplementary Figure 4). Applying the same rationale between F_ST_ and divergence to our LguyP-LguyS, where F_ST_ = 0.158, indicates that the the derived population (LguyP) separated from the ancestral sylvatic population (LguyS) approximately 330 years ago (95% CI 1041-1703 CE). Given the stochastic nature of coalescent processes between sites in genomes, the difference between the kDNA locus and nuclear genome divergence estimates is unsurprising and we do observe a large variance in nucleotide diversity along the genome as a consequence (see Figure 3D), as expected from population genetics theory (41, 42). As the formation and urbanisation of Manaus took place between 1600 and 1800 (43), these two estimates indicate that the LguyP-LguyS separation is probably within the time scale of European settlement in the region.

Given the relatively recent emergence of the LguyP, the newer population may be adapting to a new vector species and/or a new reservoir. To look for genomic signatures of adaptation as recent selective sweeps, we employed a combination of three orthogonal genome diversity metrics; local F_ST_ to measure genetic distance between populations; Tajima’s D to identify differential departures in the allele frequencies within populations relative to neutrality and the cross-population extended haplotype homozygosity (XP-EHH) test to identify extended stretches of homozygosity in one population compared to another. All these tests were applied to genomic windows to detect signals consistent with recent selection in one or both populations (Figure 3 E and F).

We found evidence for ongoing selective sweeps in two genomic regions, which had significant XP-EHH p-values, high F_ST_ and low Tajima’s D (Figure 3 E and F). These regions co-localized with runs of lower nucleotide diversity in LguyP compared to LguyS, consistent with selective sweeps in the newer city-proximal population (Figure 3 E and F, Supplementary Figure 11, 12 and 13). The selective sweeps in chromosome 15 and 34 have respectively 11 (4 hypothetical) and 68 (28 hypothetical) genes, which hampered the identification of the biological functions of genes in these regions (Supplementary Table 2). This was an exploratory analysis that should be re-evaluated when more L. guyanensis isolates become available, and when experimental analyses identify the biological function of these potential causal genes is characterized.

Collectively, these findings indicate that LguyP and LguyS populations diverged after human settlement in the Manaus region, with the derived LguyP population now occupying rural and transitional areas, and the ancestral LguyS population remaining in more distant heavily covered forested regions. We also observe that there is little genetic exchange between these species, and we see some preliminary evidence for selection between populations. This is in agreement with a potential presence of different vectors and/or reservoirs closer to the city, leading to distinct transmission cycles in these two locations.

## Discussion

This is the first large-scale population genomics characterization of *L. guyanensis* in the intersection between the Brazilian Amazon rainforest, and its largest human settlement, the city of Manaus. We revealed signs of a recent emergence of the population LguyP, city proximal, from the sylvatic population LguyS. The distinct urban proximity and forest coverage patterns observed around the residences of patients infected with parasites from each population, together with our approximate divergence estimates is suggestive of an anthropogenic environmental change driven separation and adaptation of these two *L. guyanensis* populations.

*L. guyanensis* is a member of the *Leishmania* Viannia clade, which is endemic to South America (44), and is among the major causative agents of cutaneous leishmaniasis (CL) in the Brazilian Amazon (23, 45) (22, 46). Population genomic studies within the *L. Viannia* subgenus have primarily focused on *L. braziliensis* and *L. peruviana*, revealing geographically structured populations and frequent hybridization events in regions where species ranges overlap. (19–21). A recent study of *L. braziliensis* concluded that ∼35% of genetic variation could be explained by environmental and geographic factors, indicating the importance of environmental change to the spread of leishmaniasis throughout history (19). However, the dynamics of CL micro-foci transmission between sylvatic and urban environments remain poorly understood.

We identified differences in deforestation patterns in the vicinity of residences of patients infected with the two *Leishmania guyanensis* populations, with evidence suggesting that these changes were human-driven. In the last decades, Manaus urbanisation was marked with inadequate urban planning, often associated with poverty and poor sanitary conditions for residents (47–53). These conditions likely facilitated the establishment of CL transmission cycles along the urban periphery, where habitats differ from those of the sylvatic cycle. A potential geographic isolation and adaptation to new vectors/reservoirs is in accordance with the low inter-population recombination and the presence of selective sweeps in LguyS. As obligate parasites, *Leishmania* species must co-evolve when vectors or and/or reservoirs change environments, or when they adapt to new reservoirs. Evidence from other zoonotic parasites shows similar patterns. Farmland expansion, new settlements and slums create sylvatic-urban transition zones bringing people, vectors, and reservoirs into closer contact, increasing disease transmission. Such transitions have been associated with disease emergence for other zoonoses with wildlife reservoirs, such as yellow fever, Nipah virus, Lyme disease, cholera and African trypanosomiasis (54–57). In these cases, anthropogenic land use drastically altered the composition of wildlife communities, increasing the abundance of vector/reservoir species that are better adapted to survive in urban areas (57–62). Seasonal floods in Manaus are increasing in frequency and severity (63, 64), and these may also isolate vectors or reservoirs and contribute to the dynamics of *Leishmania* populations.

The transmission dynamics of *L. guyanensis* and the other *Leishmania* species in the Manaus regions is complex, and not well understood. Although the primary vector and reservoirs of *L. guyanensis* are likely to be respectively *Nyssomya umbratilis* and *Choloepus didactylus* (two-toed sloth), *L. guyanensis* can also be found in other vectors, such as *Lutzomyia anduzei* and *Lutzomyia whitmani* (25–28). *L. guyanensis* can also infect a wide range of mammals, such as *Tamandua tetradactyla* (southern tamandua), *Proechimys sp*. (spiny rat) and *Didelphis marsupialis* (opossum) (*25, 26, 28–32*). In addition, several populations of the vector *Nyssomyia umbratilis* are known to circulate around Manaus (65, 66), and these populations have differences in their competence to transmit *L. guyanensis* (*67*), and their other aspects of their biology (68) that could affect their distribution. Deforestation is likely to alter interactions between sand flies and wild fauna, increasing the frequency of blood feeding in humans, causing an increase in the number of CL cases (69).

Our analysis was based on parasites that were isolated from human patients, so is unable to provide mechanistic or quantitative detail on the vector/reservoir cycle of CL in this region. Further research into the parasites, vectors and reservoirs in Manaus and in other human settlements in the Amazon is crucial to understand the impact of urbanization in the spread of *Leishmania*. Our expectation is that such research will lead to the identification of key targets for CL environmental management at urban/sylvatic interfaces.

## Material and methods

### Sample collection, genome sequencing

A collection of 77 *Leishmania* primary isolates from human patients with cutaneous leishmaniasis (CL) from Prof. Jorge Guerra’s repository at the Fundação de Medicina Tropical Heitor Vieira Dourado (FMTHVD) in Manaus were used in this work. Collections and the subsequent analysis were covered by these ethical approvals; CAEE: 63489722.5.0000.0068; CAEE: 80862417.3.0000.0005; University of York Biology Ethics Committee: DJ202212 and DJ202412. The access to these *Leishmania* strains as genetic heritage followed Brazilian biodiversity legislation and was registered at Sisgen (Sisgen AE85886). All *Leishmania* strains collected for this work were obtained in routine diagnostic biopsies from cutaneous lesions from patients that resided in the city of Manaus or neighbouring regions in the Amazon Rainforest, and were admitted and treated at the “Ambulatório de Dermatopatias de pacientes com Leishmaniose” from Fundação de Medicina Tropical Dr. Heitor Vieira Dourado (FMT HVD). Diagnostic protocol included the use of PCR to identify the *Leishmania* species that was infecting the patient (22, 34, 35). The cryopreserved parasites were briefly cultured in Schneider’s media with 10% foetal bovine serum, at 26°C, until they reached a density of 10^8^ parasites/mL. The parasites were centrifuged at ∼1,500g, washed once with PBS and re-suspended in DNA/RNA Shield (Zymo Research). Parasite DNA was extracted using the QIAGEN Blood and Tissue kit (QIAGEN) following manufacturer’s instructions, and sequenced at Macrogen (Seoul, South Korea), using the Nextera XT DNA library preparation kit and Illumina NovaSeq sequencer, generating 151 nucleotide long paired end whole genome sequencing (WGS) read libraries. The raw read libraries were deposited in the European Nucleotide Archive under Study Accession XXXXXX [in progress, to be completed prior to publication].

To support the classification of the Amazonian *Leishmania* isolates, representative WGS read libraries from the main *Leishmania Viannia* species that causes CL in the Amazon region were obtained from NCBI SRA, using Fastq-dump (https://github.com/ncbi/sra-tools). These correspond to 30 *L. peruviana*; 24 *L. panamensis*; 19 *L. braziliensis*; 5 *L. guyanensis*; 3 *L. naiffi*; 2 *L. lainsoni* and 1 *L. shawi* samples (Supplementary Table 1).

### Read mapping and SNP calling

To remove low quality reads and adapter sequences, WGS reads from both the Amazonian dataset as well as from the NCBI SRA dataset were trimmed using fastp v.0.23.2 (70), removing reads with quality below Phred 20, shorter than 50 nucleotides and removing bases in the N and C terminal with quality below Phred 20, where only reads where both pairs were above these cutoff values were used in downstream analysis. These libraries were mapped to the *L. braziliensis* MHOM/BR/75/M2904 2019, version 64 (TriTrypDB - (71, 72)), which contains both the nuclear chromosomes as well as the mitochondrial chromosome (kDNA - Maxicircle) using BWA mem v.0.7.17 (73). PCR duplicates and reads with mapping quality lower than 20 were removed using samtools v.1.17. Read group names were assigned with Picard tools (https://broadinstitute.github.io/picard/).

For each sample, SNPs and indels were called using Freebayes v.1.3.6 (https://github.com/ekg/freebayes), accepting calls with a minimum alternative allele read count ≥ 5; and GATK HaplotypeCaller v.4.4 (74, 75); only accepting calls discovered by both programs. *Leishmania* isolates with more than 5% of missing data, as well as SNP positions with more than 20% of missing data were excluded using VCFtools v.0.1.16 (76). The multi-sample VCF was filtered to contain only biallelic sites, with SNP quality > 1000, Minor Allele Count (MAC) of at least 1 (remove non-variant sites), read depth higher than ¼ and lower than 4x the mean coverage of all SNP sites; and mapping quality 30 on both reference and alternate alleles using BCFtools v.1.15.1 (77) (bcftools view -m 2 -M 2 -i ’MAC >=1 & QUAL > 1000 & INFO/DP > 1652 & INFO/DP < 26447 & INFO/MQM >30 & INFO/MQMR >30); and to remove insertion/deletions using VCFtools v.0.1.16 (77). This resulted in 980,755 segregating sites (SNP sites that vary at least once in the evaluated population).

For the kDNA SNPs, the SNP calling was the same as described for the nuclear genome, but restricting the region analysed to the non-repetitive kDNA region (positions 170 - 16,414) and using less stringent filtering parameters (bcftools view -m 2 -M 2 -i ’MAC >=1 & QUAL > 30 & INFO/MQM >30 & INFO/MQMR >30); and we removed insertion/deletions using VCFtools v.0.1.16 (77). This resulted in a total of 27 segregating sites.

### Complexity estimation and gene/chromosome copy number variation

The chromosome copy number variation and genome coverage was estimated using the median read depth coverage of genes in a given chromosome with non-outlier coverage (Grubb’s tests, with P<0·05), normalised by the genome coverage, estimated by the coverage of all single copy genes in the genome, using Samtools depth (78), as described in (37). The heatmap representing the CCNV was generated using R using the library pheatmap (https://github.com/raivokolde/pheatmap).

To evaluate if any of the 77 primary isolates from the Amazon region have evidence of being multiclonal infections and/or polyploid, we estimated their Complexity Index, as described in (36), using previously generated estimations of genome coverage and CCNV. To that end, the multi-sample VCF was separated into single-sample VCFs, removing non-variant sites using VCFtools v.0.1.16 (--non-ref-ac-any 1) (76) and submitted to the pipeline described in (https://github.com/jaumlrc/Complex-Infections).

### Phylogeny and population structure

For the PCA, Admixture and phylogeny analysis, SNPs with strong linkage disequilibrium (*r*^2^ > 0.5) were removed using PLINK v1.90b7 (79, 80), resulting in 409,711 segregating sites. To generate the nuclear genome phylogeny network for all *Leishmania* samples, the unlinked SNPs multisample VCF was converted to the nexus format using vcf2phylip (https://github.com/edgardomortiz/vcf2phylip), which was used as input for the NeighborNet method in SplitsTree V4.19.2 (81). The phylogeny network plot was generated using ggtree (82). Population structure analysis using the unlinked SNPs multisample VCF was performed with Admixture v1.3, which was run with K=1-20. Values of K=5 and K=10 were selected based on Cross Validation (CV) analysis.

Principal Component Analysis for the sets: 1-Whole genome, including Amazon primary isolates and NCBI samples; 2-Whole genome with only Amazon primary isolates; and 3-Individual chromosomes with only Amazon primary isolates; was generated with PLINK (79, 80). Both Admixture and PCA plots were generated in R, using ggplot (83).

The kDNA phylogeny was estimated by maximum likelihood, using IQ-TREE2 (84), with automatic selection of the best nucleotide substitution model (Best-fit model: HKY+F chosen according to BIC) and 1,000 bootstrap replicates. The tree plot was generated in ITOL (85). To estimate the mean kDNA SNP counts for LguyP and LguyS as well as the kDNA *d*_XY_; consensus kDNA fasta sequences were generated for each *L. guyanensis* samples, incorporating their specific SNPs using BCFtools consensus (77, 86). These sequences were combined in a single multi-fasta file, and the estimation of within and between mean kDNA SNP counts for LguyP and LguyS as well as the kDNA *d*_XY_ were performed in R, using the packages seqinR (87), ape (88) and adegenet (89).

### Introgression analysis

To detect and visualise introgression patterns between LguyP and LguyS populations, we performed Diagnostic Index Expectation Maximisation (DIEM) analysis using the diemr package v1.4.3 in R (39) (https://cran.r-project.org/web/packages/diemr/). DIEM identifies diagnostic SNPs that distinguish two populations and calculates hybrid indices representing the proportion of ancestry from each population. VCF files containing biallelic SNPs from 71 *L. guyanensis* samples were converted to DIEM format. To maximise population resolution, we employed a whole-genome approach rather than per-chromosome analysis, as the latter yielded insufficient SNPs per chromosome for reliable introgression detection. Population assignments were defined using LguyP (n=51) and LguyS (n=10) as reference populations, with 10 samples of uncertain assignment retained in the analysis but excluded from reference population definitions. Analysis parameters included: markerPolarity = FALSE (allowing the expectation-maximisation algorithm to determine optimal allele polarities) and uniform diploid ploidy specification. Diagnostic SNPs were filtered at the 60th percentile of diagnostic index (DI) values, retaining 15,962 markers. Genotypes were polarised and hybrid indices calculated for all samples. Chromosome paintings display individuals ordered by hybrid index, with colours representing ancestry: red (homozygous for LguyP-associated alleles), blue (homozygous for LguyS-associated alleles), and yellow (heterozygous).

### Sequence diversity, recombination and selective sweep analysis

The nucleotide diversity (π) was estimated for 1-Each evaluated *Leishmania* species with 10 or more samples (*L. braziliensis*, *L. guyanensis*, *L. panamensis* and *L. peruviana*); and 2- *L. guyanensis* sub-populations LguyP and LguyS individually (see section: **Phylogeny and population structure**, and the results section), in 10 kb windows using VCFtools (76). Tajima’s D (10 kb windows) and F_ST_ were estimated for the *L. guyanensis* sub-populations LguyP and LguyS, using VCFtools (76). The statistical difference of the minor allele frequency and π between LguyP and LguyS estimated using the Mann-Whitney U test, in R, with a significance value of p < 0.05. The presence of ongoing and recent selective sweep events between populations LguyP and LguyS was assessed with the R package *rehh* (90), using the Cross Population Extended Haplotype Homozygosity (XP-EHH) test (91).

To identify SNPs that are enriched in one of the Amazonian *L. guyanensis* populations (LguyP and LguyS) the multi-sample VCF was loaded in R with vcfR (92) and the data was separated into LguyP and LguyS. Then, the Alternate Allele Frequency (AAF) for each population was estimated by dividing the Allele Count (AC) by the total number of available alleles (i.e 102 (51*2, as it is diploid); for LguyP and 38 (19*2 as it is diploid) for LguyS). Then, for each SNP position, the LguyP AC was subtracted from the LguyS AC, generating the “AC difference” (ACD). SNP positions with ACD higher than 0.5 were classified as having a higher occurrence in a given population.

### Evaluation of the forest cover and deforestation in the local of residence of patients infected with LguyP or LguyS

To assess the potential association between forest habitat and different *L. guyanensis* populations, the sample points around the residence of patients from whom the *Leishmania* isolates were obtained were compared in terms of the forest cover and age of recent forest disturbance. A map of percentage forest cover derived from Landsat satellite imagery at a resolution of 30 m by 30 m pixels (93) was used to assess the percentage forest cover in the 3 km radius regions (buffer zones) around each sample point. Mann Whitney U tests of difference were then applied to assess differences in forest cover between samples from different *L. guyanensis* populations. Data showing the year of deforestation since 2000, released along with the forest cover map (93), was considered using the same methods. It allowed the investigation of the difference in age of forest disturbance between population samples. Buffer zones with different radius were also investigated from 1 km through to 10 km distances, where the 3 km radius represents a commonly walkable distance around localities (c. 30 minutes walking time). The figures were generated in R (v4.4.2 or above)(94, 95).

## Acknowledgements

This work was supported by the UK:Brazil Joint Centre Partnership in Leishmaniasis, which was funded by the UK Medical Research Council/Fundação de Amparo a Pesquisa do Estado de Sao Paulo (JCPiL)(MR/S019472/1). JLRC and DCJ were also supported by the JCPiL and by a MRC New Investigator Research Grant to DCJ (MR/T016019/1). The University of York Viking high performance compute facility was used during this project. We are grateful for computational support from the University of York, IT Services and the Research IT team. HG acknowledges the support from Conselho Nacional de Desenvolvimento Científico e Tecnológico (CNPq; research fellowship to H.G. #308000/2023-4). We acknowledge the technical support of Denison Vital de Jesus (in memoriam). We acknowledge the support from Fundação de Amparo a Pesquisa do Estado de Sao Paulo (FAPESP fellowship 2019/24393-1 to LCR) .

## Supplementary figures

**Supplementary figure 1:**
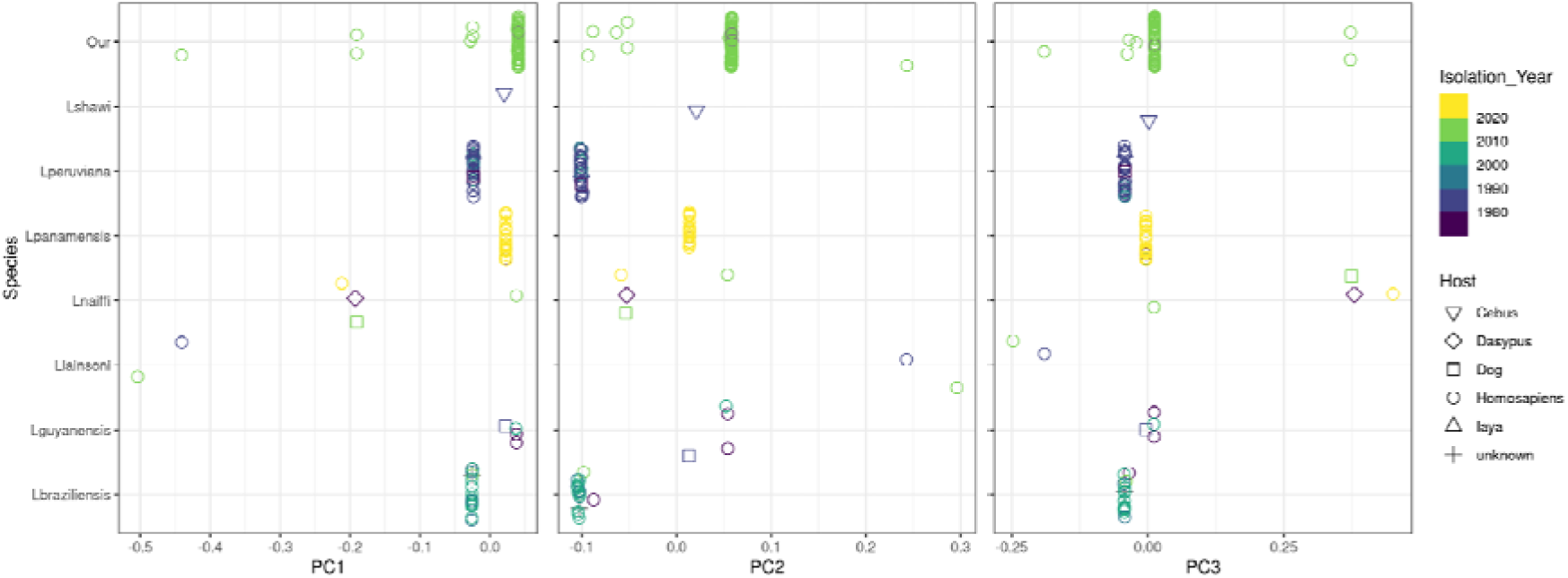
PCA from the WGS SNPs confirms the phylogeny network classification of the *Leishmania* amazonian samples. Each panel corresponds to a different principal component (PC), while the X axis represents the PC coordinate. The Y axis represents each species collection. The year of isolation of each sample is represented in colours, while the host of isolation is represented by shapes.

**Supplementary Figure 2:**
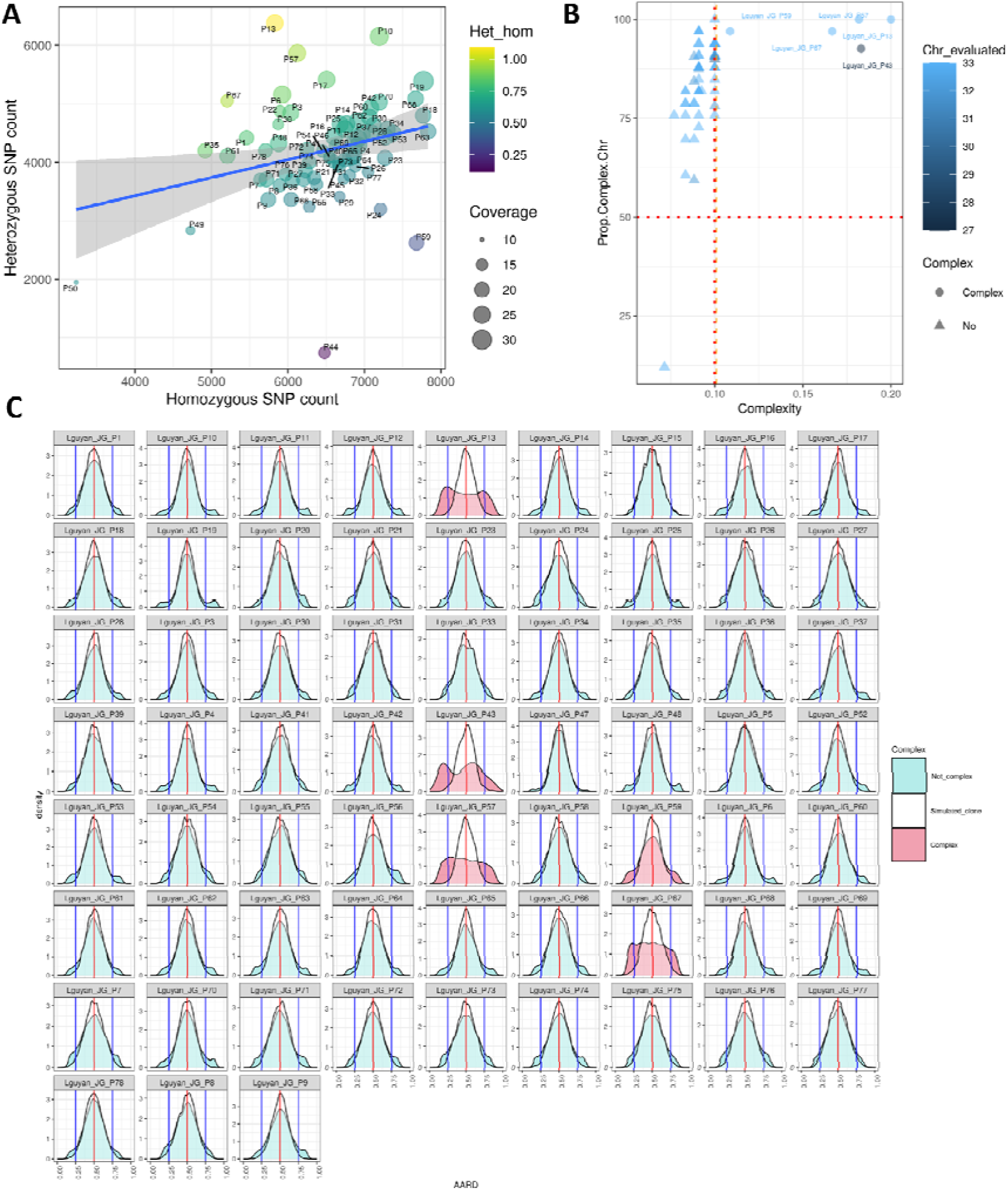
*L. guyanensis* primary isolates heterozygosity and Complexity. **A)** Assessment of Heterozygous (Y axis) and Homozygous (X axis) in *L. guyanensis* primary isolates. The ratio of heterozygous/Homozygous SNPs is represented by a colour scale, from low (blue) to high (yellow) values. The genome coverage is represented by size. **B)** Complexity estimation on 66 *L. guyanensis* and other primary isolates classified as L. guyanensis in PCR, that had genome coverage > 15x and at least 100 heterozygous SNPs. **C)** Density distributions of the AARD proportion in heterozygous positions. Each panel corresponds to a different primary isolate, and the colours red, blue and white correspond respectively to complex, non-complex and the simulated diploid clonal data.

**Supplementary Figure 3:**
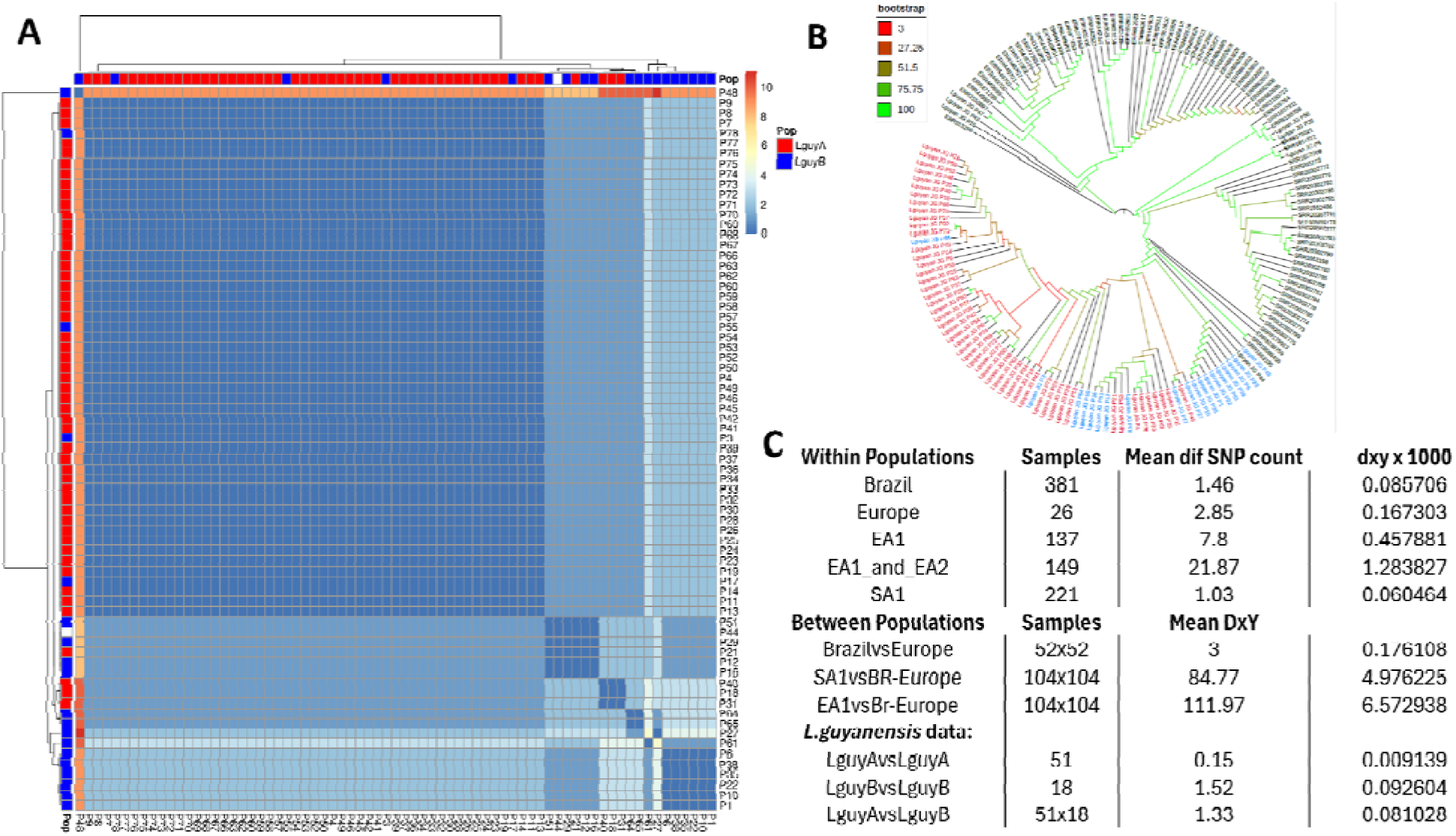
SNP evaluation in the kDNA (mitochondrial DNA) of *L. guyanensis*. **A)** Heatmap showing the number of kDNA SNPs between each pair of L. guyanensis samples. The samples were reordered based on the manhattan distance of their average distances (UPGMA). The colorstrip colour corresponds to the population of origin of each sample, red for LguyP and blue for LguyS. **B**) Maximum likelihood phylogeny of the kDNA sequence of all Leishmania Viannia samples. **C)** Table showing the “Mean number of different SNPs between isolates from LguyP and LguyS (Mean dif SNP count) and their DxY.

**Supplementary Figure 4:**
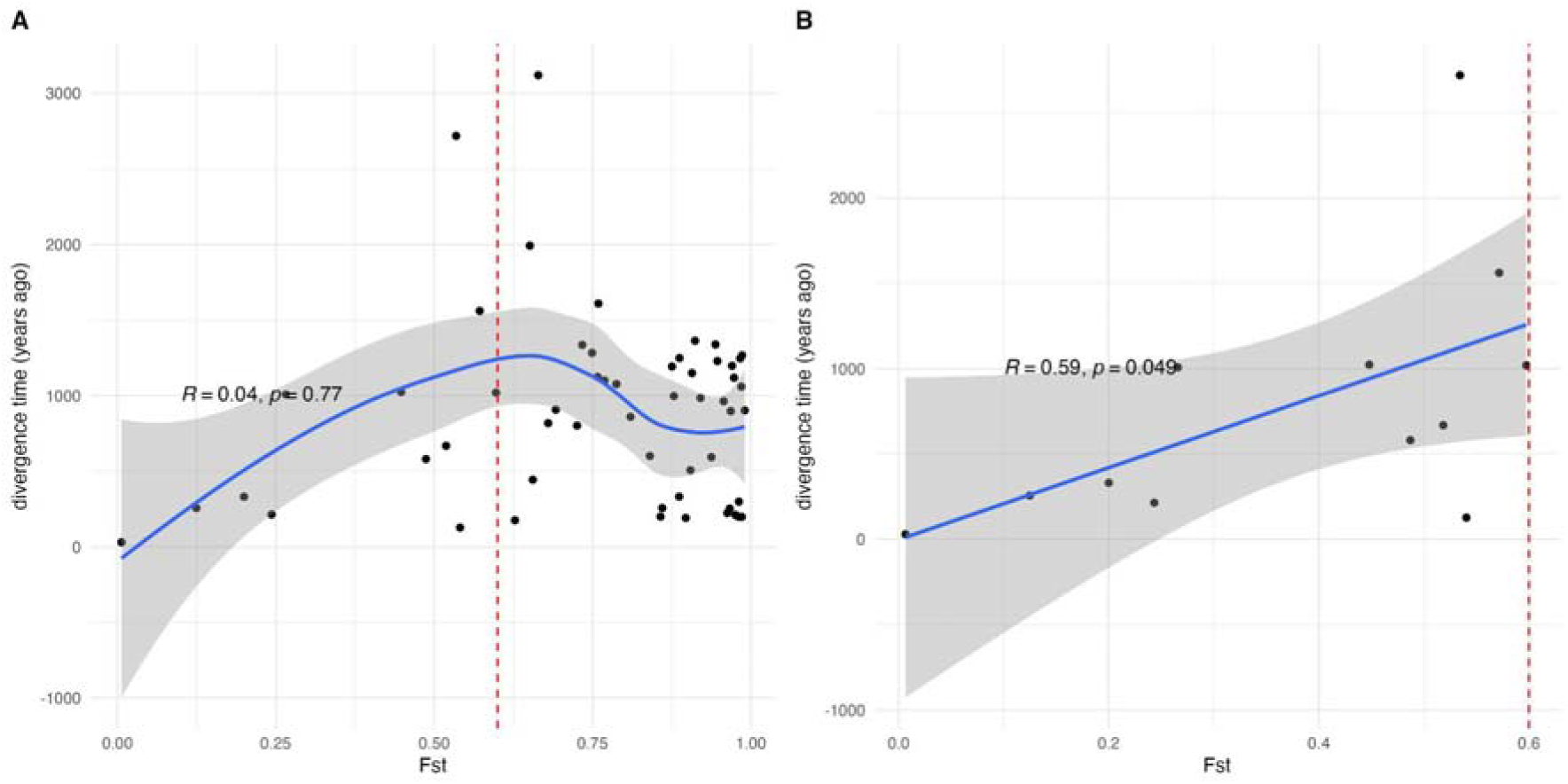
Population divergence time estimation using F_ST_. In our previous analysis of *Leishmania donovani* species complex divergence [1], we showed that genetic distances measured using F_ST_ were correlated with population divergence times measured from kinetoplastid (mitochondrial) using coalescent time divergence dating estimated using BEAST [2]. To obtain a very approximate estimate of the divergence time between the *L. guyanensis* populations LguyP and LguyS, we assume the same neutral divergence with time. From *L. donovani* data, we show **A)** BEAST split dates (years ago) appears to correlate with F_ST_ up to approximately F_ST_ = 0.6, when F_ST_ is likely saturated. In **B)** we show the BEAST split dates (years ago) relationship with F_ST_ for values ≥ 0.6. In both panels the blue line shows the regression, a loess moving regression for panel A and a linear model for panel B, with the grey shading showing regression 95% confidence intervals. Spearman rank correlations are indicated. This is not significant for all F_ST_ values (R = 0.04, P = 0.77), but is significant for F_ST_ values ≤ 0.6 (R = 0.59, P = 0.049). The linear model for panel B intercepts with the 0,0 point (F_ST_ = 0, years ago = 0), as we would expect for one non-diverged panmictic population. if we apply this relationship to *L. guyanensis,* the linear model for F_ST_ ≤ 0.6 values predicts that a population pair with F_ST_ = 0.1578 we observed for the *L. guyanensis* populations divided 330 years ago, in 1689 (330 years before the mean *L. guyanensis* sample collection date of 2019). The 95% confidence interval is 984 to -322 years ago, ie: 1035 CE to the present date, since -322 years ago is nonsensical.

**Supplementary Figure 5:**
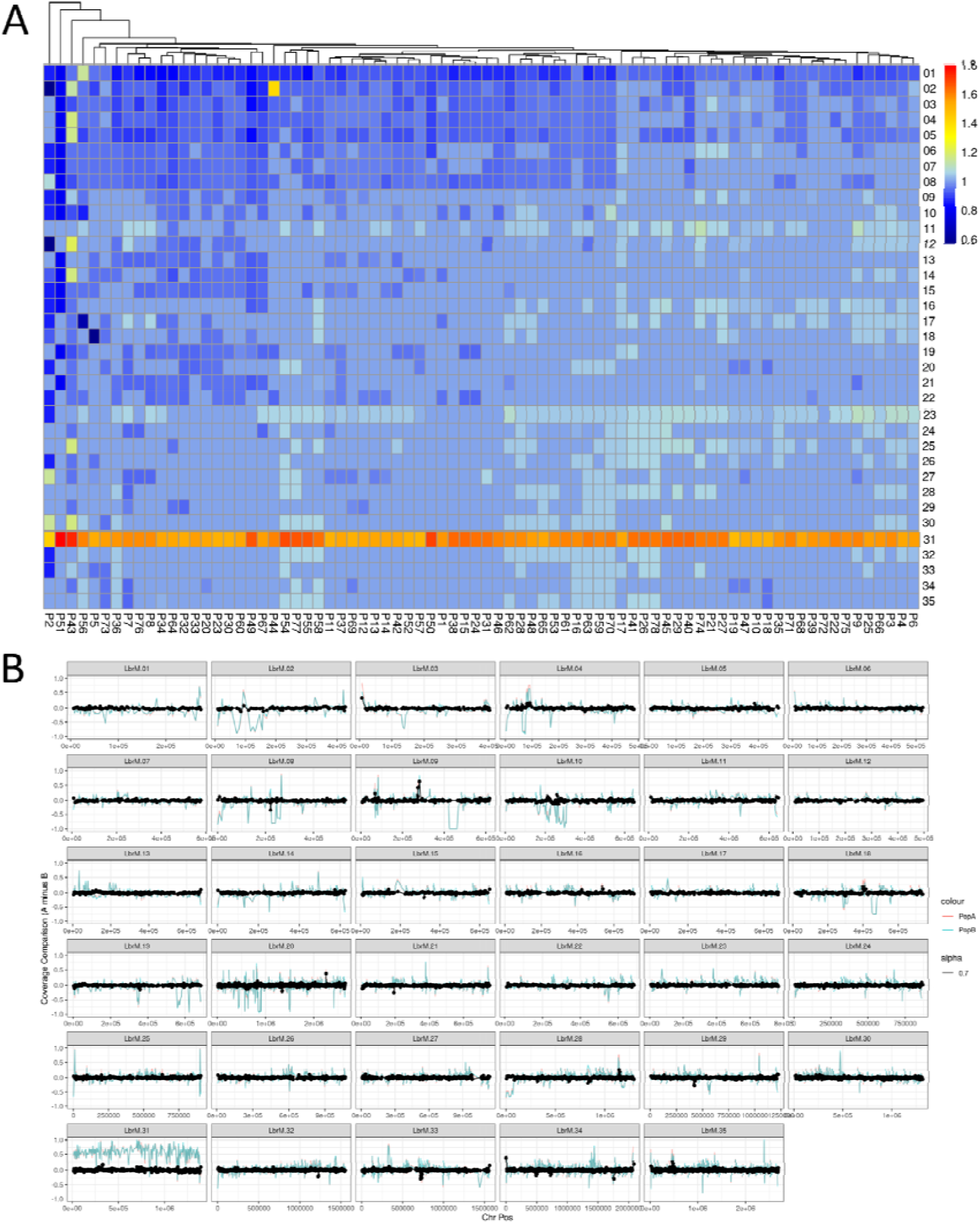
Chromosomal and gene copy number variation in the Leishmania primary isolates from the Amazon. **A)** Heatmap representing the chromosomal copy number variation (CCNV) in all samples. Each line represents a different chromosome, while each column represent a different isolate. The chromosome copy number is represented by colours, ranging from blue (low) to red (high) copies. B) Comparison of gene copy numbers between LguyP (red) and LguyS (blue). Each panel corresponds to a different chromosome, where the X axis corresponds to the chromosome position. The black dots correspond to the difference in the gene copy number between the mean coverage in LguyP and LguyS. The blue and red lines represent the gene copy number in LguyP or LguyS subtracted by 1.

**Supplementary Figure 6:**
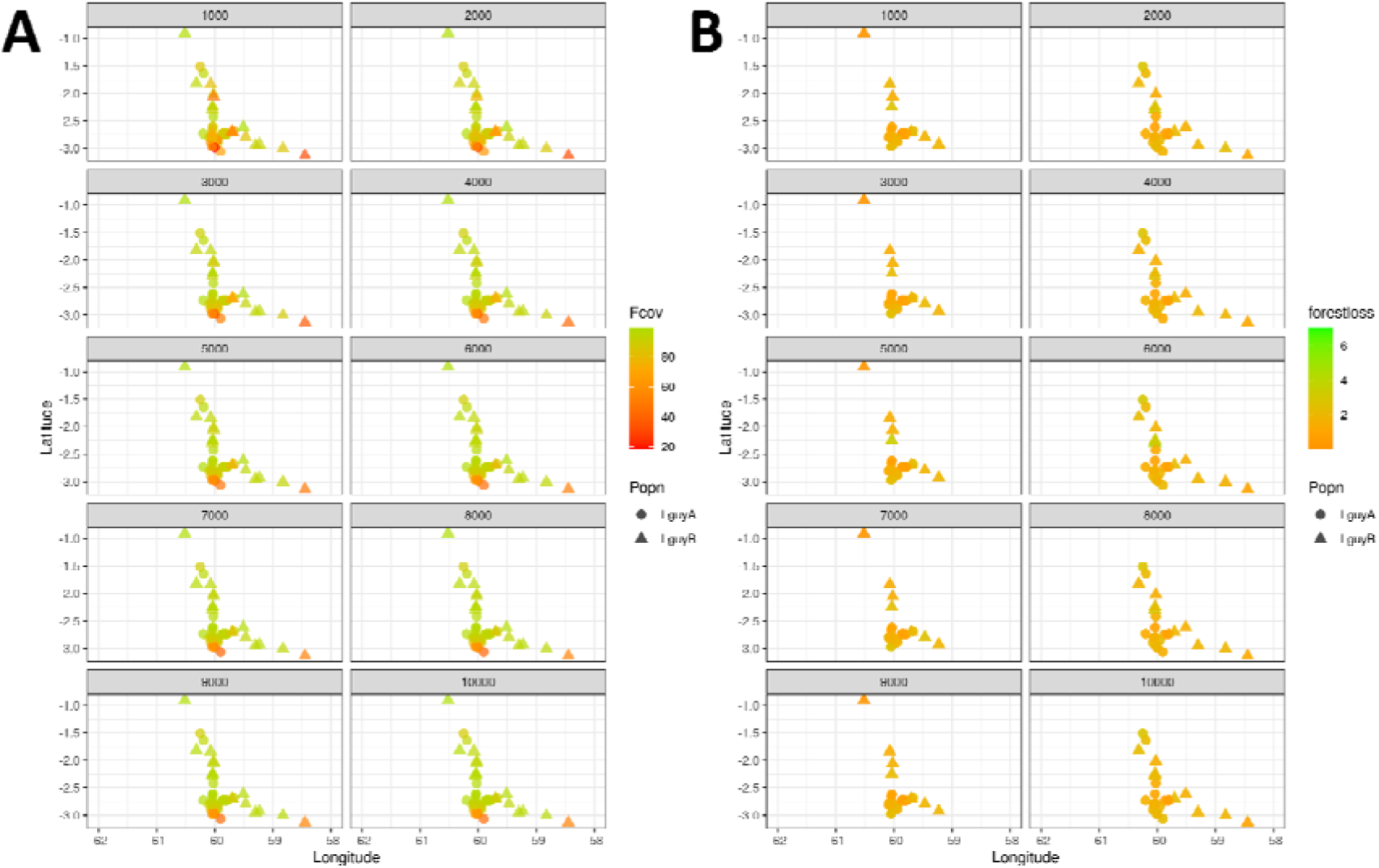
Forest coverage and forest loss after the year 2000 around the residence of the patients infected with *L. guyanensis*. **A)** Forest coverage and **B**) forest loss after the year 2000 around the residence of the patients infected with *L. guyanensis*. Each panel corresponds to a different radius around the residence, varying from 1000m to 10,000m.

**Supplementary Figure 7:**
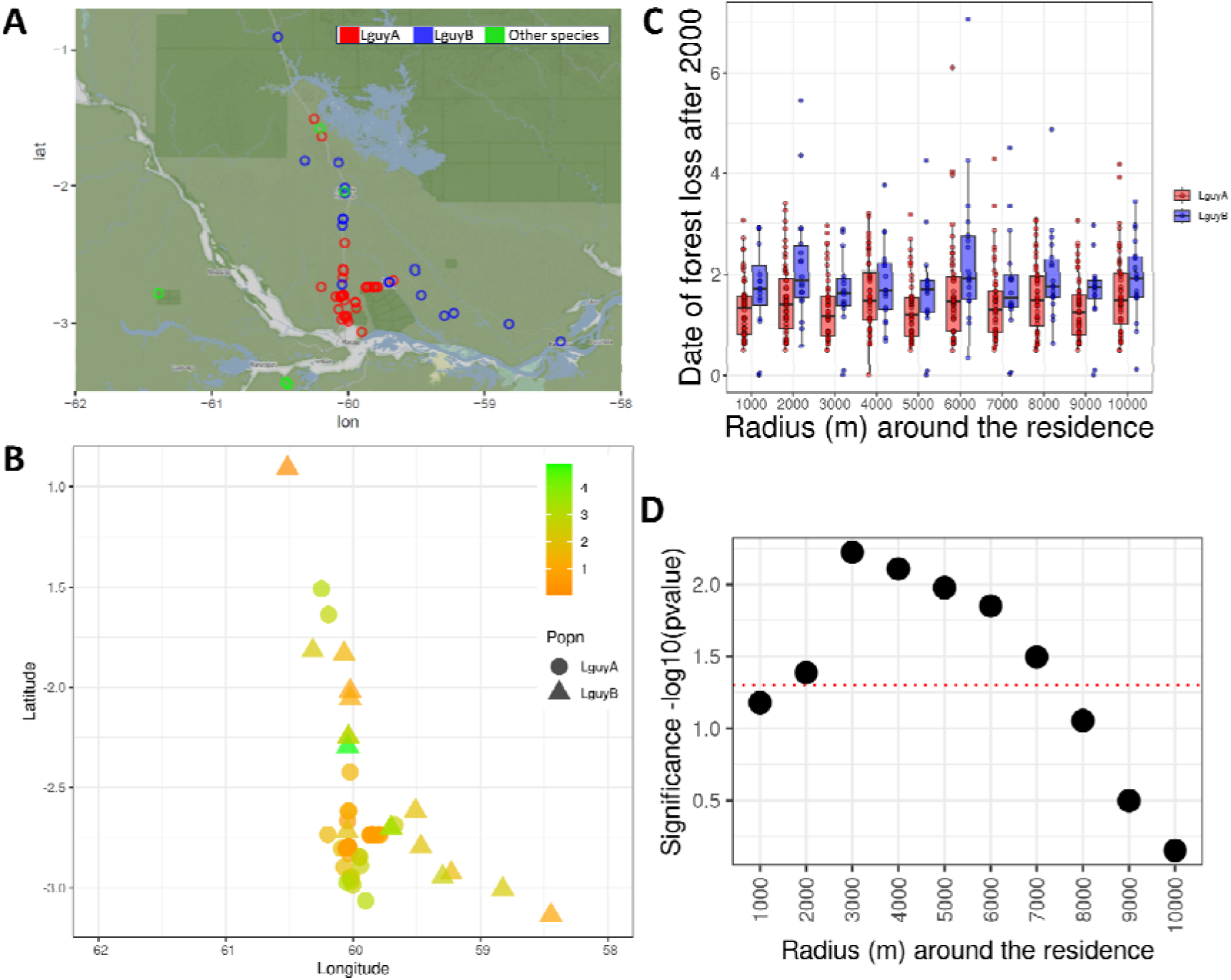
Forest loss after the year 2000 around the residence of the patients infected with *L. guyanensis*. **A)** Geographic distribution of the locality of residence of the patients from whom each *L. guyanensis* primary isolate was obtained. **B)** Number of years after the year 2,000 that the deforestation occurs. Lower values correspond to older deforestation, while large values correspond to recent deforestation. **C)** Box Plots representing the number of years after the year 2,000 that the deforestation around the local of residence of patients occurred, with increasing radius values (1000 to 10,000 m). **D)** P-values for the different radius shown in “C”. The red line corresponds to the significance value of 0.05.

**Supplementary Figure 8:**
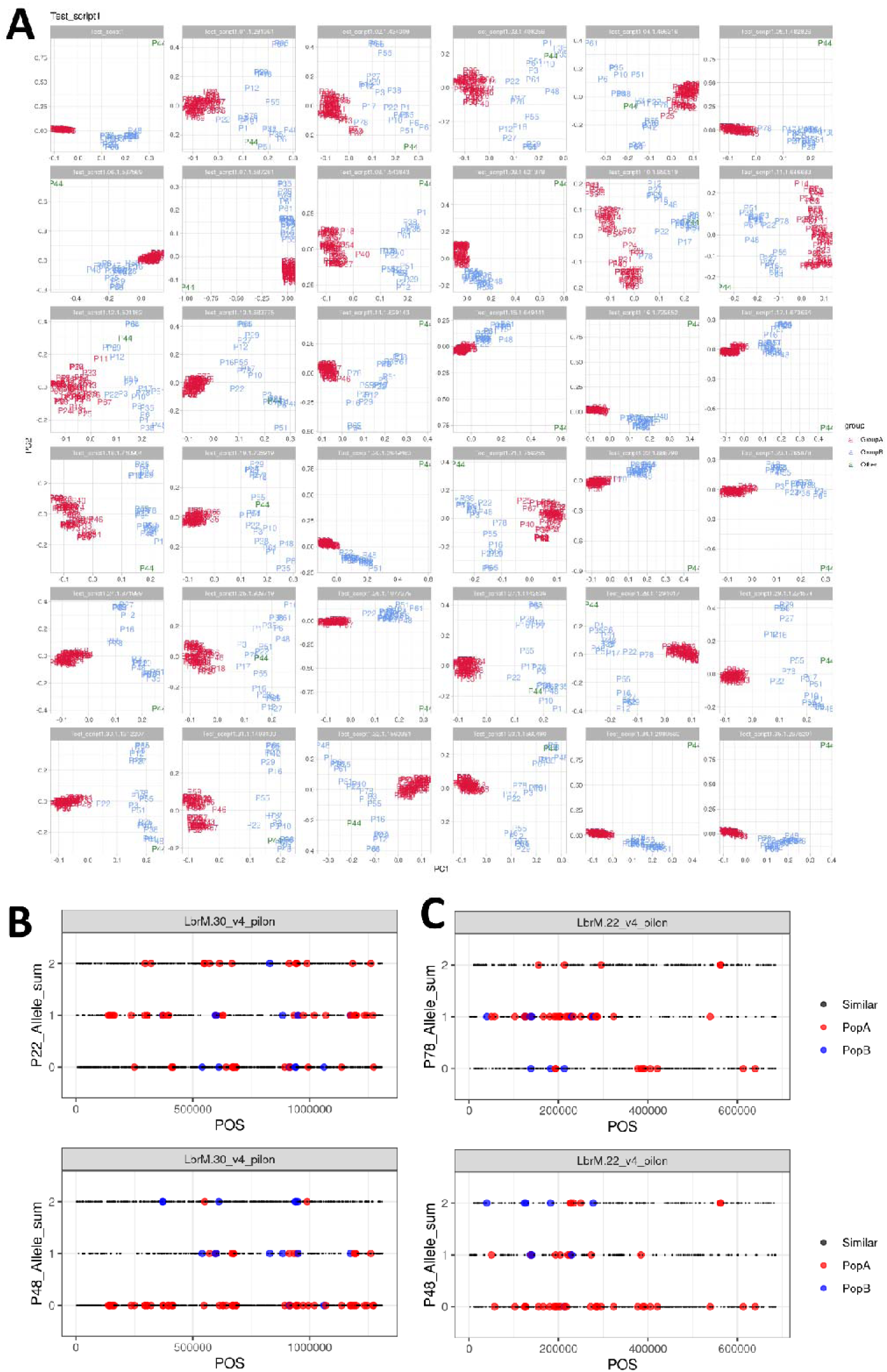
Chromosome SNP PCA and SNP genotypes supports LguyP and LguyS population, but shows potential recombination events. **A)** Each panel corresponds to a different chromosome, with the exception of the first panel, that has the result for the whole genome, similar to the Figure 2A. LguyP and LguyS isolates are respectively coloured in red or blue. Only two strains appeared to be intermediate between populations from this PCA analysis in one chromosome each; P22 in chromosome 30 and P78 in chromosome 22. Both these strains generally cluster with LguyS (blue), but appear closer to LguyP (red) in these genomic regions. **B,C)** To examine the possibility that this pattern is due to recent genetic exchange, we examined SNP genotypes within these strains in these chromosomes, focusing on the few ‘population diagnostic’ variant positions where the allele frequency of the LguyP and LguyS populations differ by ≥ 0.5 (ie: 0.3 in one population and 0.8 in another). We ‘paint’ the population diagnostic SNPs that are enriched in LguyS with red dots, and LguyS enriched SNPs with blue dots (consistent with PCA colouring). All other SNPs are indicated with smaller black dots. In these plots the *y* axis corresponds to the allele count in the strain considered: 0 indicates homozygous at the diagnostic allele, 1 indicates heterozygous state, and 2 the other homozygous state. The *x* axis corresponds to the chromosome position. **Part B) upper panel** shows SNP genotypes in P22 in chromosome 30, with LguyP-enriched alleles (red) and LguyS-enriched alleles (blue). **Part B) lower panel** shows strain P48, whose chromosomes always cluster with LguyS (blue). We observe that P22 has more LguyS (red) alleles in heterozygous and homozygous states than the non-introgressed strain P48. **Part C) upper panel** shows SNP genotypes in P78 in chromosome 22 painted as above, and for comparison in **part C) lower panel we show** SNP genotypes in from the LguyP strain P48 in chromosome 22. Again we observe that the proposed introgressed strain P78 has more LguyS (red) alleles in heterozygous and homozygous states than the non-introgressed strain P48. Both introgressed strains P22 and P78 have an excess of heterozygous SNPs, consistent with an F1 hybrid or similarly early recent genetic change.

**Supplementary Figure 9:**
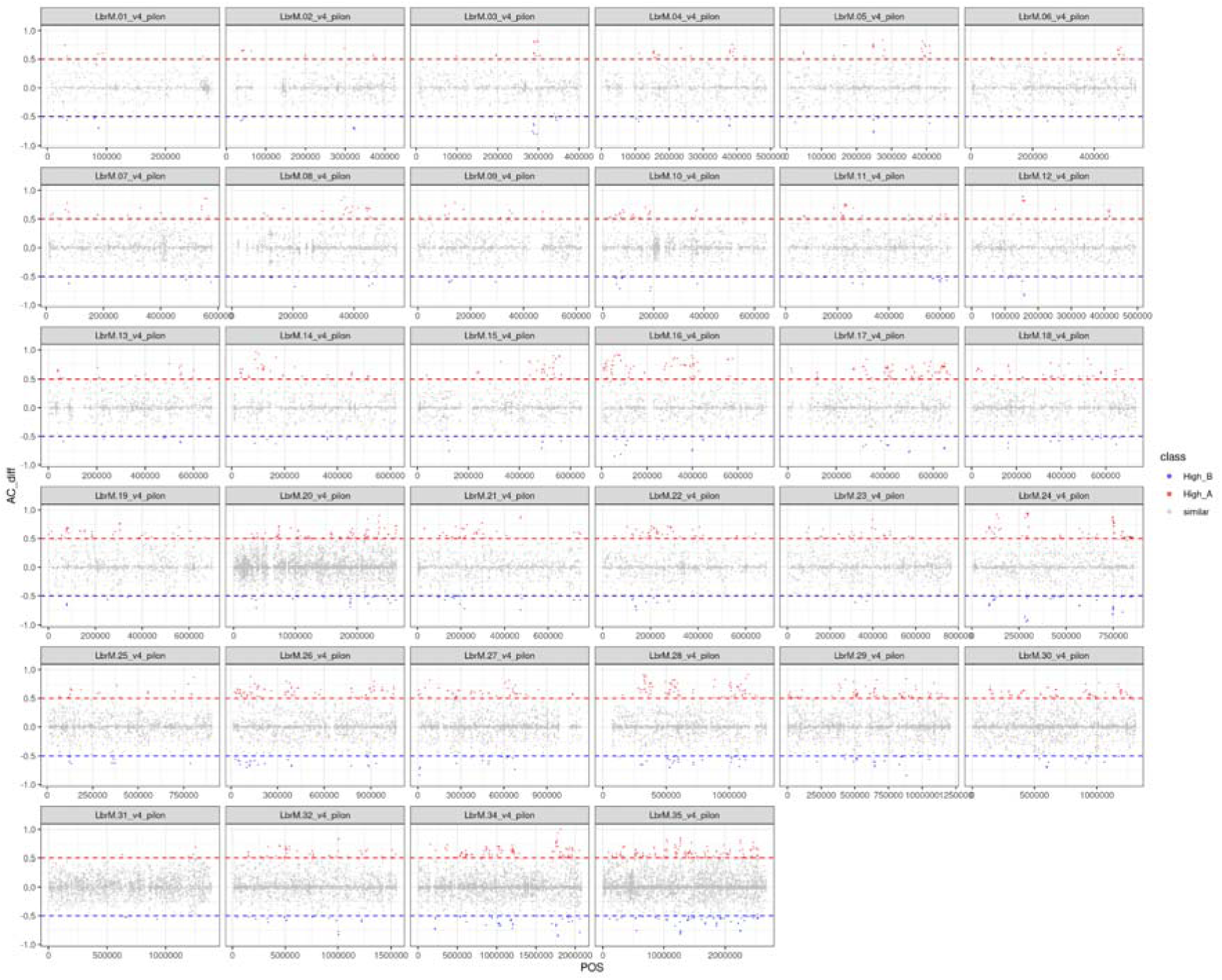
SNPs that are enriched in each population. In this plot, each panel corresponds to a chromosome, where the X axis represents the chromosome position. The Y axis represents the normalised Allele Count (AC) difference between LguyP and LguyS. SNPs that are enriched in LguyP are in red, while the ones enriched in LguyS are in blue. SNPs that have a similar occurrence in both are represented in gray.

**Supplementary Figure 10:**
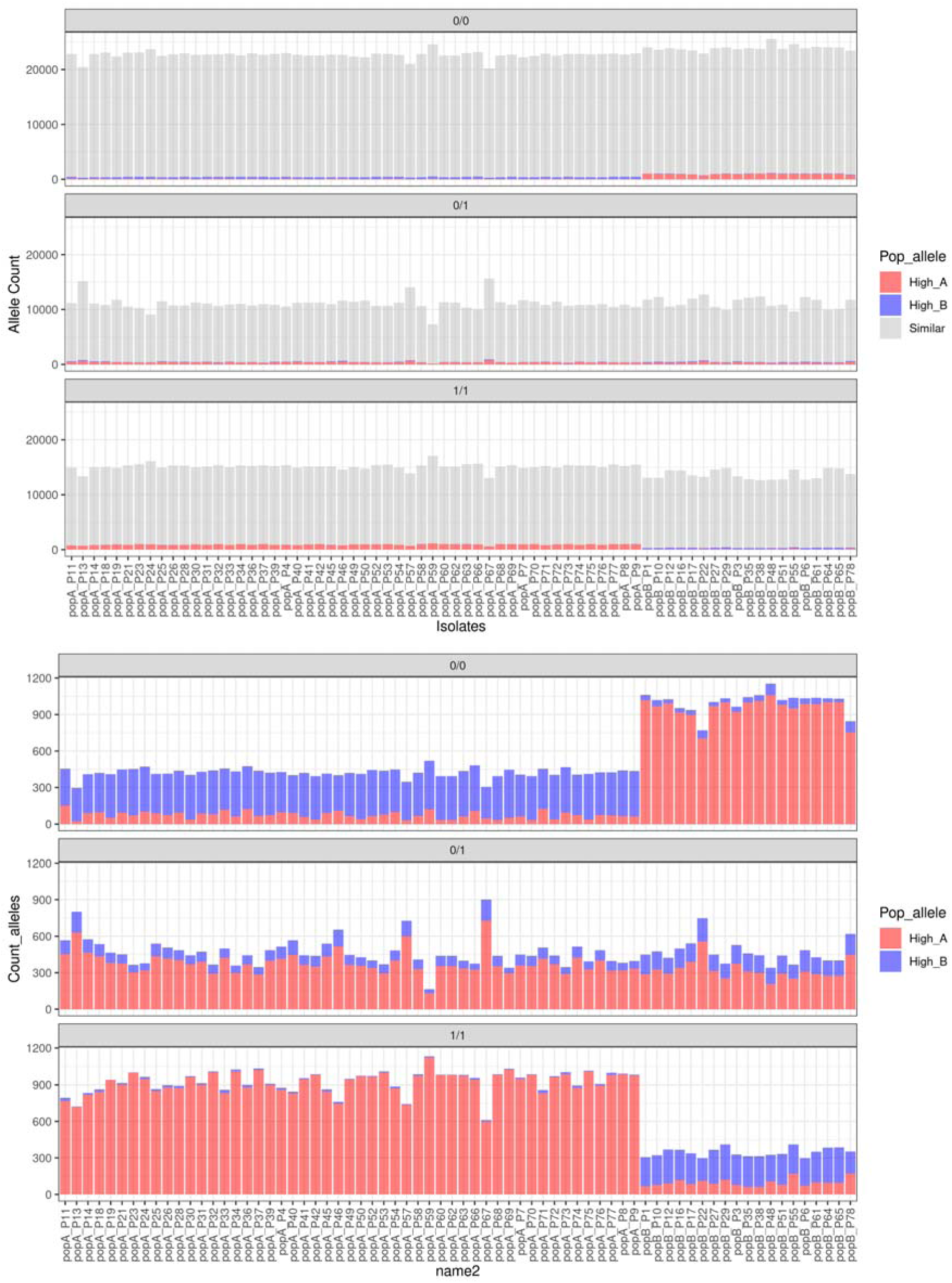
Correspondence between the genotypes in each isolate with the primary genotype in LguyP or LguyS in segregating sites in the population. Each panel corresponds to a different genotype: 0/0 Reference homozygous; 0/1 Heterozygous; 1/1 Homozygous alternate allele. The colour represents the “enrichment” of the alternate allele in LguyP (red), LguyS (blue) or similar in both (grey). The high of the bar corresponds to the number of SNPs observed in each class. This means that for isolates from LguyP, we expect high numbers of 1/1 that are red and high numbers of 0/0 that are blue; while for LguyS we expect high levels of 1/1 that are blue and high levels of 0/0 that are red. Deviations of this trend might mean recombination. All 19 LguyS isolates are on the right of the graph. **A)** Including the SNPs that have a similar occurrence in LguyP and LguyS. **B)** Excluding the SNPs that have a similar occurrence in LguyP and LguyS

**Supplementary figure 11:**
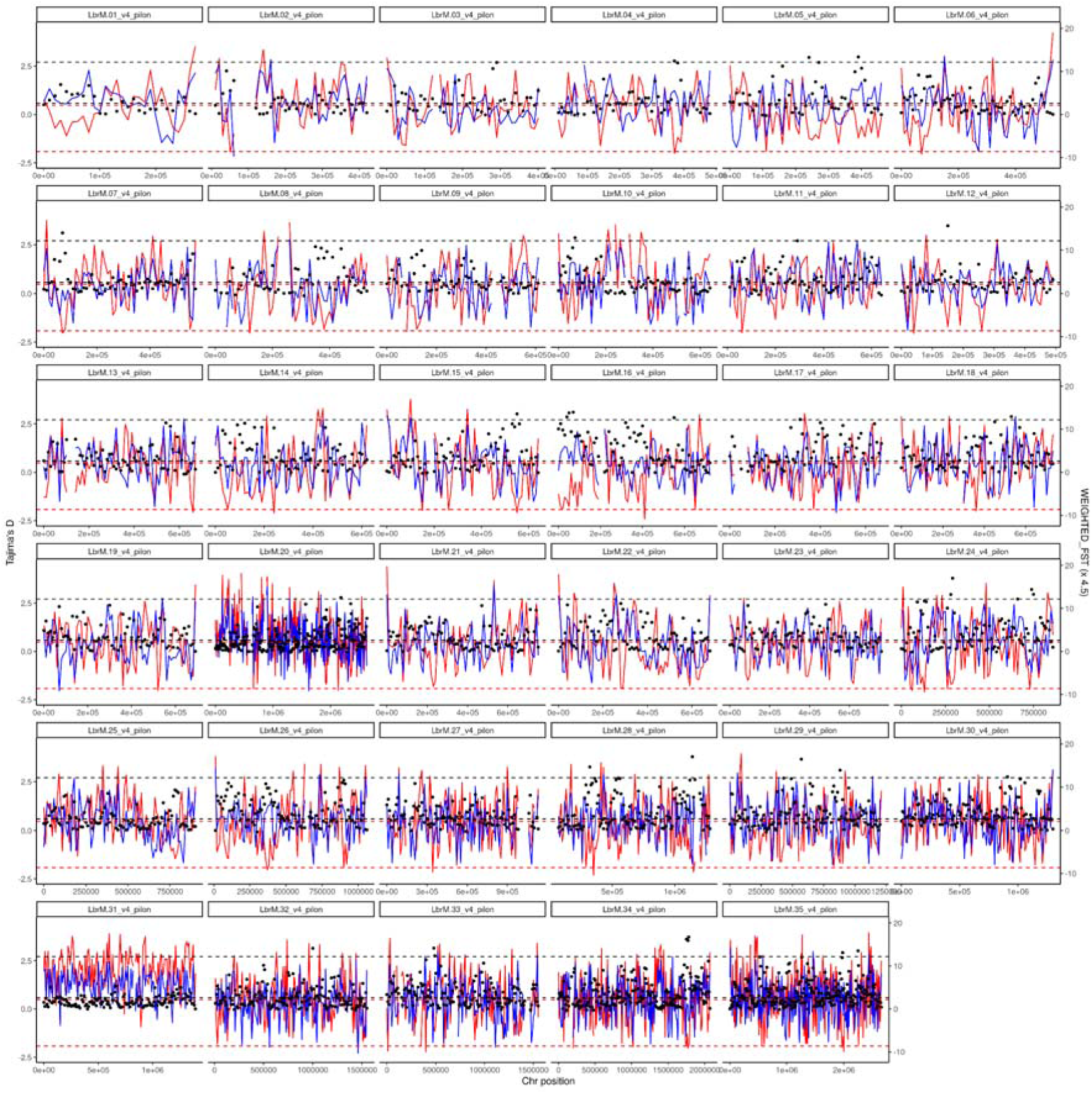
FST and Tajima’s D value for each chromosome in LguyP and LguyS. Each panel corresponds to a different chromosome. Left Y axis represents the scale for the Tajima’s D estimation, where each population is represented by coloured lines: red: LguyP; blue: LguyS; and FST: right Y axis, black dots along the chromosomal region.

**Supplementary Figure 12:**
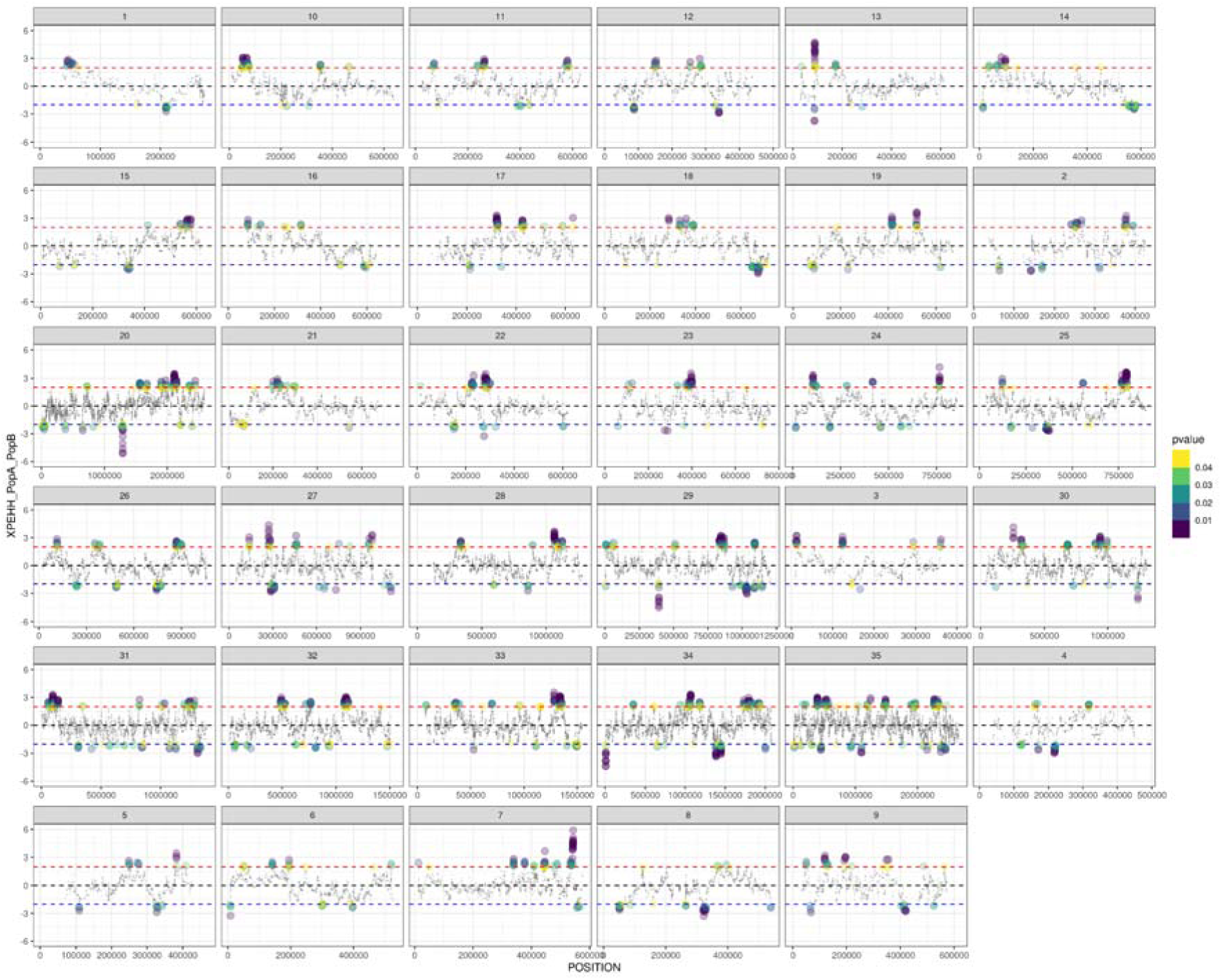
XP-EHH results (Y axis) for each SNP position along the chromosome comparing LguyP and LguyS. Each panel corresponds to a different chromosome. Each dot corresponds to one SNP, where significant p-values are larger and coloured. Values above the red line correspond to potential selective sweeps on LguyP, while values below the blue line to potential sweeps in LguyS.

**Supplementary Figure 13:**
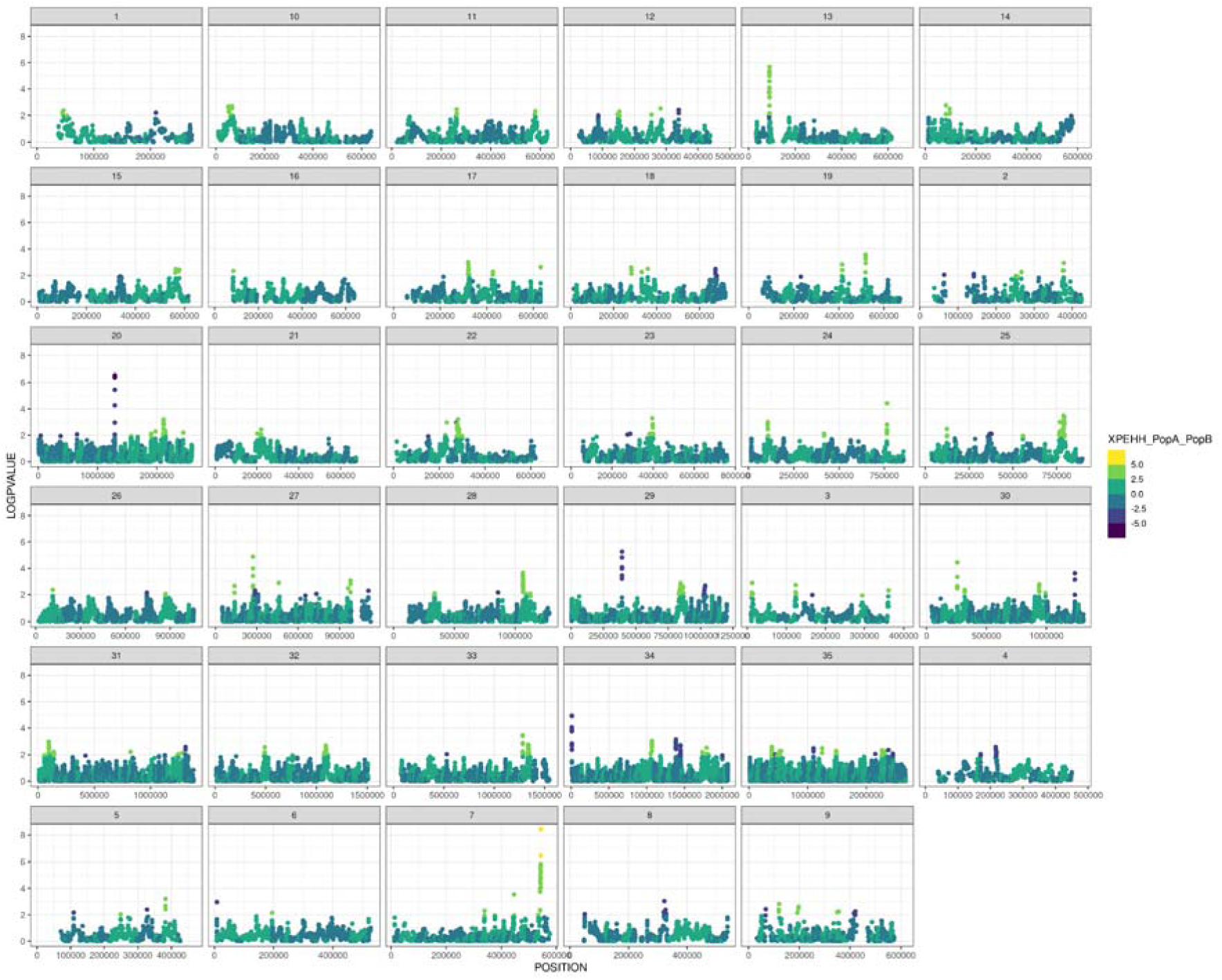
XP-EHH log(p-values) (Y axis) for each SNP position along the chromosome comparing LguyP and LguyS.

**Supplementary Figure 14:**
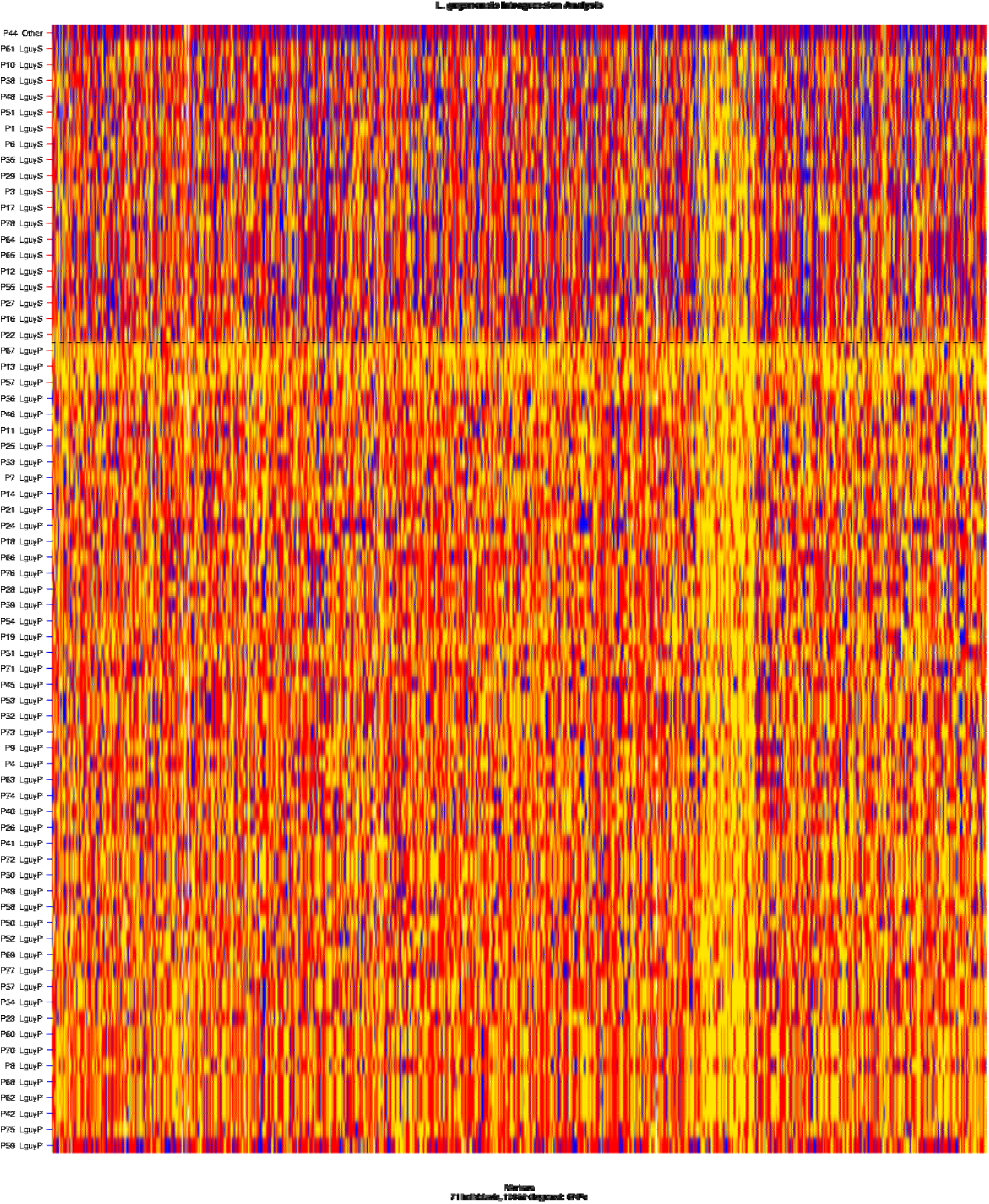
Diagnostic Index Expectation Maximisation polarisation of markers. Chromosome painting showing marker polarisation with SNPs from the 71 *L. guyanensis* samples analysed using Diagnostic Index Expectation Maximisation (*diem*). Each row represents an individual sample ordered by hybrid index determined by *diem* (proportion of LguyS ancestry), with samples arranged from highest (top) to lowest (bottom) values. LguyS (city-proximal) samples occupy the upper portion of the plot (indicated by the blue bar), whilst LguyP (sylvatic) samples occupy the lower portion (indicated by the red bar). The dashed horizontal line indicates the transition between populations. Each column represents one of 15,962 diagnostic SNP markers filtered at the 60th percentile of diagnostic index values. Colours indicate ancestry state: blue = homozygous for LguyS-associated alleles, red = homozygous for LguyP-associated alleles, yellow = heterozygous, white = missing data. Regions of LguyP (red) ancestry in predominantly LguyS samples indicate gene flow (LguyP to LguyS). The LguyS sample P22 that appears to be closer to LguyP strains in the PCA analysis of chromosome 30 (Figure 3A), does not appear to contain the large LguyP ancestry blocks (red) that we would expect from recent gene flow.

